# EEF2-inactivating toxins engage the NLRP1 inflammasome and promote epithelial barrier disruption upon *Pseudomonas* infection

**DOI:** 10.1101/2023.01.16.524164

**Authors:** Miriam Pinilla, Raoul Mazars, Romain Vergé, Leana Gorse, Karin Santoni, Kim Samirah Robinson, Gee Ann Toh, Laure Prouvensier, Stephen Adonai LeonIcaza, Audrey Hessel, David Péricat, Marlène Murris, Anthony Henras, Julien Buyck, Céline Cougoule, Emmanuel Ravet, Franklin L. Zhong, Rémi Planès, Etienne Meunier

**Author notes:** (RP), (EM). Institute of Pharmacology and Structural Biology (IPBS), University of Toulouse, CNRS, Toulouse, France. These authors equally supervised this work (Rémi Planès, Etienne Meunier).

## Abstract

The intracellular inflammasome complex have been implicated in the maladaptive tissue damage and inflammation observed in chronic *Pseudomonas aeruginosa* infection. Human airway and corneal epithelial cells, which are critically altered during chronic infections mediated by *P. aeruginosa*, specifically express the inflammasome sensor NLRP1. Here, together with a companion study, we report that the NLRP1 inflammasome detects Exotoxin A (EXOA), a ribotoxin released by *P. aeruginosa* Type 2 Secretion System (T2SS) during chronic infection. Mechanistically, EXOA-driven Eukaryotic Elongation Factor 2 (EEF2) ribosylation and covalent inactivation promotes ribotoxic stress and subsequent NLRP1 inflammasome activation, a process shared with other EEF2-inactivating toxins, Diphtheria Toxin and Cholix Toxin. Biochemically, irreversible EEF2 inactivation triggers ribosome stress-associated kinases ZAKα- and P38-dependent NLRP1 phosphorylation and subsequent proteasome-driven functional degradation. Finally, Cystic Fibrosis cells from patients exhibit exacerbated P38 activity and hypersensitivity to EXOA-induced ribotoxic stress-dependent NLRP1 inflammasome activation, a process inhibited by the use of ZAKα inhibitors. Altogether, our results show the importance of *P. aeruginosa* virulence factor EXOA at promoting NLRP1-dependent epithelial damage and identify ZAKα as a critical sensor of virulence-inactivated EEF2.

**KEY POINTS:**

- *P. aeruginosa* induces NLRP1-dependent pyroptosis in human corneal and nasal epithelial cells
- *P. aeruginosa* Exotoxin A (EXOA) and other EEF2-inactivating bacterial exotoxins activate the human NLRP1 inflammasome
- EEF2 inactivation promotes ribotoxic stress response and ZAKα kinase-dependent NLRP1 inflammasome activation.
- Bronchial epithelial cells from Cystic Fibrosis patients show extreme sensitivity to ribotoxic stress-dependent NLRP1 inflammasome activation in response to Exotoxin A
- P38 and ZAKα inhibition protects Cystic Fibrosis epithelial cell from EXOA-induced pyroptosis

## INTRODUCTION

*Pseudomonas aeruginosa* is an opportunistic bacterial pathogen that can cause acute and chronic life threatening infections (Qin et al., 2022; Maurice et al., 2018). Due to widespread antibiotic resistance and its adaptation to the airway, skin and cornea of immune-compromised patients (e.g. ciliated dyskinesia, chronic granulomatous diseases, cystic fibrosis), *Pseudomonas aeruginosa* is listed as an important ESKAPE pathogen in the WHO list of microbes of concerns (De Oliveira et al., 2020). *P. aeruginosa* can trigger acute infections thanks to the expression of a Type 3 secretion system (T3SS), which leads to a robust inflammatory reaction mostly mediated by monocytes/macrophages and neutrophils (Qin et al., 2022). However, during chronic infections *P. aeruginosa* switches into a metabolically different state that represses T3SS expression and allows the expression and secretion of a different arsenal of effectors involved in the formation/maintenance of biofilm-like structures, which are extremely resistant to antibiotic treatments (Qin et al., 2022). In addition, the secretion of numerous matrix remodeling factors such as (phospho)lipases, proteases, siderophores, oxidative and toxic molecules strongly contribute to immune response deviation as well as to tissue damages, including epithelial barrier disruption (Qin et al., 2022). In this context, inflammatory mediator analysis from Cystic Fibrosis (CF) patients chronically infected with *P. aeruginosa* highlights an enrichment in inflammasome-derived cytokine IL-1β, suggesting that during the chronic step of *P. aeruginosa* infection one or many inflammasomes might be activated (Bonfield et al., 2012). Given the prominent epithelial cell damage observed in *P. aeruginosa* infected patients, and the poorly addressed function of the epithelial barrier in antibacterial defense, we hypothesized that some epithelial inflammasomes might respond to one or various factors released by *P. aeruginosa* during chronic infections.

Inflammasomes, which mostly belong to the Nod-Like Receptor (NLR) and AIM2-Like Receptor (ALRs) families, are a subset of germline-encoded innate immune sensors that detect and respond to various signs of infections and environmental stresses (Lacey and Miao, 2020). Upon activation, inflammasome-forming sensors assemble a cytosolic supramolecular structure composed of the sensor/receptor, the adaptor protein ASC and the protease Caspase-1. Inflammasome assembly leads to caspase-1-dependent maturation and release of inflammatory cytokines IL-1β and IL-18 as well as to pyroptosis, a pro-inflammatory form of cell death characterized by gasdermin (D)-driven pore formation and Ninjurin-1-dependent membrane disassembly (Kayagaki et al., 2021; Broz and Dixit, 2016; Lacey and Miao, 2020).

The human NLRP1 inflammasome is notable among other inflammasome sensors because of 1) its epithelia-restricted expression, 2) its divergence from rodent counterparts and 3) its uncommon domain organization. Rare germline mutations and single-nucleotide polymorphisms in NLRP1 are associated with infection sensitivity, skin, corneal and intestinal inflammatory disorders as well as with asthma susceptibility in humans, hence underlying an important function of NLRP1 at triggering an innate immune response in various epithelia (Zhong et al., 2016; Griswold et al., 2022).

A conserved mechanism of NLRP1 inflammasome activation relies on proteasome-driven degradation of the NLRP1 N-terminal autoinhibitory fragment (NT) and the subsequent oligomerization of the released C-terminal fragment (CT), which nucleates the NLRP1 inflammasome assembly. To this regard, recent studies identified human-specific mechanism of NLRP1 inflammasome activation, including the recognition of viral double-stranded RNA, cleavage by 3C and 3CL proteases from rhinovirus and SARS-CoV-2 viruses as well as ZAKα and P38 stress kinase-driven phosphorylation upon exposure to ribosome stressors (UV-B irradiation, anisomycin) (Tsu et al., 2021; Robinson et al., 2020; Griswold et al., 2022; Bauernfried et al., 2021; Robinson et al., 2022; Jenster et al., 2023; Fenini et al., 2018; Planès et al., 2022).

In this study, we discover that in human corneal and nasal epithelial cells, *P. aeruginosa* contributes to barrier disruption by secreting the Eukaryotic Elongation Factor 2 (EEF2)-inactivating toxin Exotoxin A (EXOA), which subsequently triggers assembly of the NLRP1 inflammasome and release of associated IL-1 cytokine as well as pyroptotic cell death. In this process, EXOA-inactivated EEF2 promotes ribotoxic stress response (RSR) and subsequent activation of the stress kinases ZAKα and P38 (Wu et al., 2020; Vind et al., 2020). Subsequently, activated ZAKα and P38 stimulate NLRP1 phosphorylation in its disordered region, functional degradation and activation. Finally, Cystic Fibrosis cells from patients show exacerbated P38 activity and hypersensitivity to EXOA-induced ribotoxic stress-dependent NLRP1 inflammasome activation, a process reverted by the use of ZAKα inhibitors. Altogether, our results describe the ability of *P. aeruginosa* virulence factor EXOA at promoting NLRP1-dependent tissue damage and identify ZAKα as a critical sensor of bacterial pathogen-driven ribosome inactivation.

## RESULTS

### *P. aeruginosa* infection activates the NLRP1 inflammasome in human corneal and airways epithelial cells

Chronic infections mediated by *P. aeruginosa* can target various sites, including skin (Spernovasilis et al., 2021), cornea (Hazlett, 2005) and airways/lung (Qin et al., 2022) tissues. In certain contexts, *P. aeruginosa* metabolically transitions into a mucoid form that enables biofilm formation. As chronically infected patients exhibit both epithelial alterations and robust IL-1β cytokine levels, we wondered about the contribution of inflammasome response from epithelial compartments during *P. aeruginosa* infection. To mimic *P. aeruginosa* biofilm-like behavior, we relied on a transwell-adapted method that allowed bacteria to grow on top of a 0.4µm porous membrane and where epithelial cells were seeded on the bottom of the wells (**Fig. 1A**). In this context, we analyzed the inflammasome response (IL-1β/IL-18 and cell death) of primary human corneal (pHCECs) and nasal epithelial cells (pHNECs) co-cultured with *P. aeruginosa*. We observed that *P. aeruginosa* triggered robust cell death and IL-1β/IL-18 release both in pHNECs and pHCECs (**Fig. 1A**). Importantly, the use of Z-YVAD, an inhibitor of Caspase-1 activity, underscored that IL-1β/IL-18 release fully depended on Caspase-1 activity while cell death was partly dependent on Caspase-1, Caspase-3 and Caspase-8 (**Fig. 1A**). This suggests that the inflammasome in pHNECs and pHCECs could contribute to IL-1β/IL-18 and cell death in response to extracellular *P. aeruginosa*. In order to determine which inflammasome is involved, we immunoblotted for various inflammasome-forming sensors in those epithelial cells (**Fig. 1B**). Although we failed at detecting NLRP3 expression, we could detect expression of the NLRP1 sensor in addition to the complete inflammasome machinery (ASC, Caspase-1, GSDMD) both in nasal and corneal epithelial cells **(Fig. 1B**) (Robinson et al., 2020; Griswold et al., 2022). Hence, we hypothesized that NLRP1 might be a sensor of extracellular *P. aeruginosa* in the airway and corneal compartments. To address this question, we used our previously described epithelial lung A549-ASC-GFP reporter cell lines in which hNLRP1 construct was stably introduced (Planès et al., 2022). Florescence microscopy and quantification of active inflammasome complexes (ASC-GFP^+^ puncta/Specks) in co-cultured reporter cells unveiled that *P. aeruginosa* exposure promoted the formation of inflammasome complexes as well as induced significant cell death specifically in NLRP1-expressing cells (**Fig. 1C**). This suggested that NLRP1 could be a sensor of a yet unknown secreted product by *P. aeruginosa*. To further determine the involvement of NLRP1 in primary cells at responding to extracellular *P. aeruginosa*, we invalidated NLRP1 expression in nasal and corneal epithelial cells by using CRISPR-Cas9 method (**Fig. 1D**). Co-culture of WT and NLRP1-deficient cells with *P. aeruginosa* showed that *NLRP1*^-/-^ cells exhibited a defect in cell death (measured by propidium iodide incorporation and LDH release) as well as at releasing IL-1β/IL-18 cytokines (**Fig. 1D, E**). As hNLRP1 inflammasome activation requires ubiquitination and subsequent proteasome-driven functional degradation, which releases the active NLRP1 C-Ter fragment, we next incubated pHCECs with PAO1 or with Val-boro-pro (Vbp), a known chemical activator of the NLRP1 inflammasome, in presence or absence of the proteasome inhibitor bortezomib (**Fig S1A-C**). Measure of the pyroptosis pore forming protein Gasdermin-D processing, hNLRP1 degradation, cell death and IL-1β release in pHCECs showed that proteasome inhibition strongly impaired those processes, hence suggesting that the hNLRP1 inflammasome activation by *P. aeruginosa* occurs in a proteasome-dependent manner in corneal and airway epithelial cells (**Fig S1A-C**).

**Figure 1.**
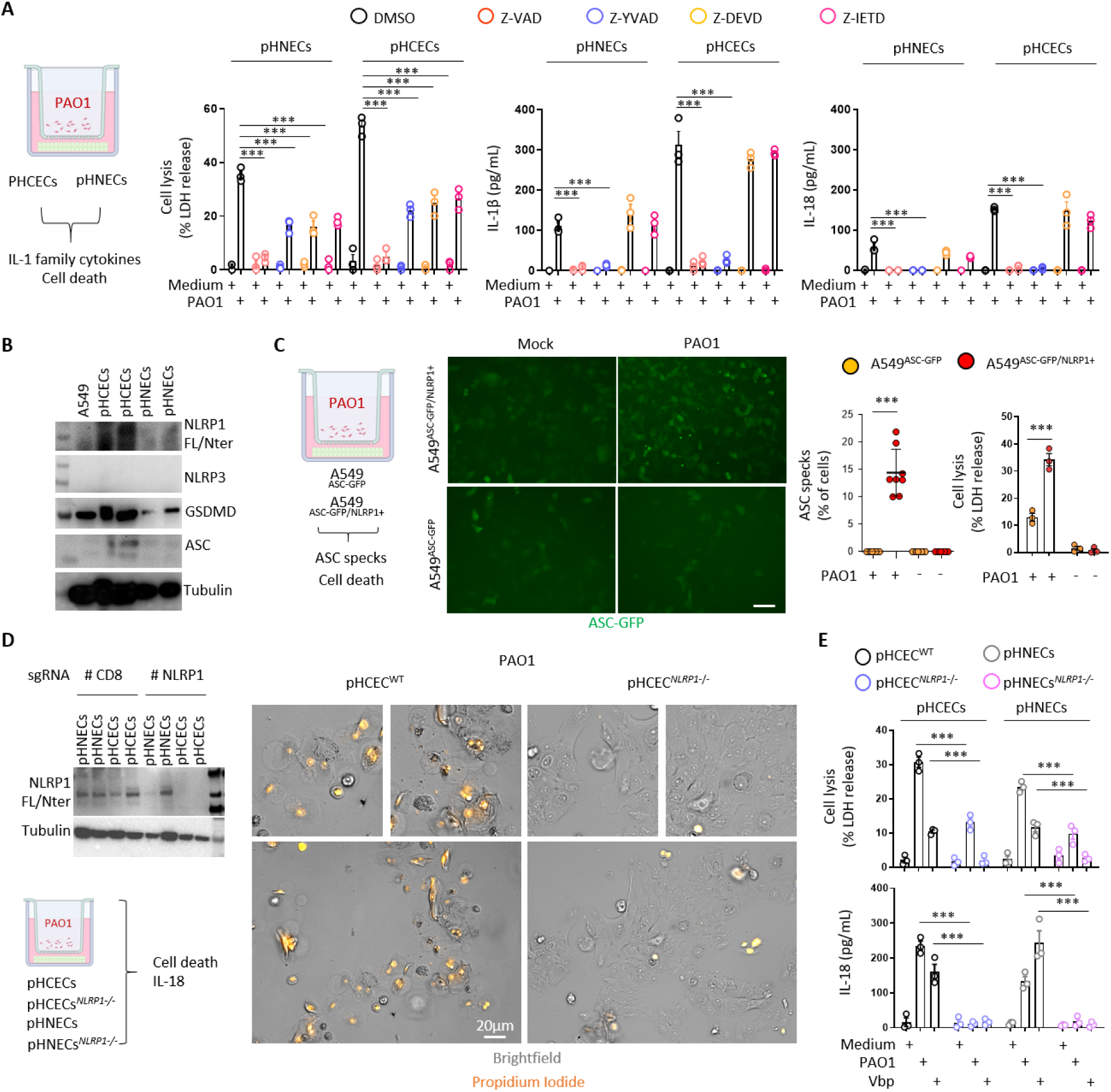
*P. aeruginosa* triggers human NLRP1 inflammasome activation in corneal and nasal epithelial cells. A. Cell lysis (LDH) and IL-1β/IL-18 release evaluation in primary Human Corneal and Nasal Epithelial Cells, respectively pHCECs and pHNECs, upon *P. aeruginosa* (PAO1, 1.10^5^ bacteria) co-culture for 24 hours. ***p ≤ 0.001, Two-Way Anova with multiple comparisons. Values are expressed as mean ± SEM. B. Immunoblotting examination of NLRP1, NLRP3, GSDMD, ASC and Tubulin in pHCECs and pHNECs. Immunoblots show lysates from one experiment performed three times. C. Florescence microscopy and associated quantifications of ASC-GFP specks in A549^NLRP1+/ASC-GFP^ and A549^NLRP1-/ASC-GFP^ reporter cell lines exposed to P. aeruginosa (PAO1, 1.10^5^ bacteria) for 24 hours. ASC-GFP (Green) pictures were taken in dish during after infection. Images shown are from one experiment and are representative of n = 8 independent experiments; scale bars, 50 µm. ASC complex percentage was performed by determining the ratios of cells positives for ASC speckles on the total cells (brightfield). At least 10 fields from each experiment were analyzed. Values are expressed as mean ± SEM. ***p ≤ 0.001, One-Way Anova. D. Immunoblotting characterization of genetic invalidation of *NLRP1* in pHCECs and pHNECs population using CRISPR-Cas9 and microscopy visualization of plasma membrane permeabilization (Propidium Iodide incorporation, orange) in pHCECs co-cultured with PAO1 (1.10^5^ bacteria) for 24 hours. Images shown are from one experiment and are representative of n = 3 independent experiments; scale bars, 20 µm. E. Cell lysis (LDH) and IL-18 release evaluation in WT or *NLRP1*-deficient pHCECs and pHNECs, upon Valbororo pro, Vbp 15µM) treatment or *P. aeruginosa* (PAO1, 1.10^5^ bacteria) co-culture for 24 hours. ***p ≤ 0.001, Two-Way Anova with multiple comparisons. Values are expressed as mean ± SEM.

### EEF2-inactivating Exotoxin A promotes NLRP1 inflammasome response

As the NLRP1 inflammasome responds to a yet to be determined secreted product of *P. aeruginosa*, we next wondered about the identity of this factor. *P. aeruginosa* expresses various secretion systems (T1SS to T6SS) that allows either secreting or injecting various elements (Filloux, 2011, 2022). Using co-cultures of A549-ASC-GFP/NLRP1 reporter cells lines with PAO1 transposon mutants for different secretion systems, we observed that Type-2 Secretion System (T2SS)-deficient PAO1 specifically failed at inducing NLRP1 inflammasome complex assembly (**Fig. 2A**). Among other factors released by PAO1 T2SS are phospholipases (PLCN, PLCH), elastase protease LASB and the Exotoxin A (EXOA) (Liao et al., 2022). Using PAO1 transposon mutants for each of those factors, we observed that only the PAO1 strain that is deficient for Exotoxin A (EXOA) lost the ability of promoting the formation of the NLRP1 inflammasome in our reporter cell lines (**Fig. 2B**). This suggests that T2SS-secreted EXOA is the major factor with which extracellular *P. aeruginosa* activates the NLRP1 inflammasome in epithelial cells.

**Figure 2.**
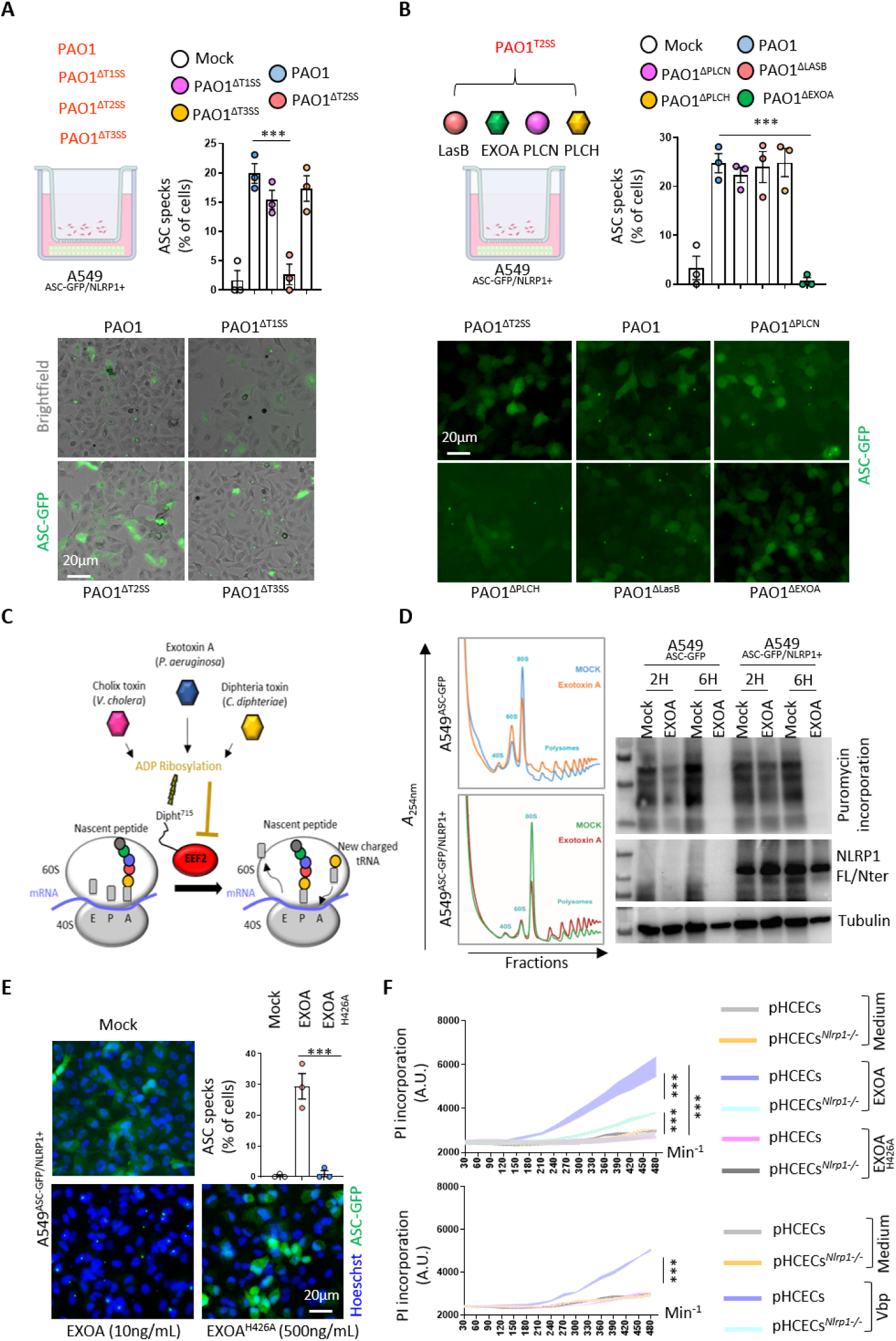
*P. aeruginosa* EEF2-inactivating Exotoxin A (EXOA) promotes NLRP1 inflammasome response. A. Florescence microscopy and associated quantifications of ASC-GFP specks in A549^NLRP1+/ASC-GFP^ reporter cell lines exposed to 1.10^5^ *P. aeruginosa* (PAO1) and associated isogenic mutants for various Secretion Systems (PAO1^ΔT3SS^, PAO1^ΔT2SS^, PAO1^ΔT1SS^) for 24 hours. ASC-GFP (Green) pictures were taken in dish during after infection. Images shown are from one experiment and are representative of n = 3 independent experiments; scale bars, 20 µm. ASC complex percentage was performed by determining the ratios of cells positives for ASC speckles on the total cells (brightfield). At least 10 fields from n = 3 independent experiments were analyzed. Values are expressed as mean ± SEM. One-Way Anova. B. Florescence microscopy and associated quantifications of ASC-GFP specks in A549^NLRP1+/ASC-GFP^ reporter cell lines exposed to 1.10^5^ *P. aeruginosa* (PAO1) and associated isogenic mutants for various Type-2 Secretion System virulence effectors (PAO1^ΔPLCN^, PAO1^ΔPLCH^, PAO1^ΔLASB^, PAO1^ΔEXOA^) for 24 hours. ASC-GFP (Green) pictures were taken in dish during after infection. Images shown are from one experiment and are representative of n = 3 independent experiments; scale bars, 20 µm. ASC complex percentage was performed by determining the ratios of cells positives for ASC speckles on the total cells (brightfield). At least 10 fields from n = 3 independent experiments were analyzed. Values are expressed as mean ± SEM. ***p ≤ 0.001, One-Way Anova. C. Schematic mechanism of *P. aeruginosa* Exotoxin A (EXOA) and related toxins at mediating EEF2 ribosylation and inactivation and subsequent ribosome inactivation. D. Determination of ribosome inactivation in A549^NLRP1+/ASC-GFP^ and A549^NLRP1-/ASC-GFP^ reporter cell lines exposed to Exotoxin A (EXOA, 10ng/mL) for 2 and 6 hours by measuring ribosome polysome accumulation and puromycin incorporation. E. Florescence microscopy and associated quantifications of ASC-GFP specks in A549^NLRP1+/ASC-GFP^ reporter cell lines exposed to EXOA (10ng/mL) or its catalytically dead mutant EXOA^H426A^ (500ng/mL) for 10 hours. ASC-GFP (Green) pictures were taken in dish during after toxin exposure. Images shown are from one experiment and are representative of n = 3 independent experiments; scale bars, 20 µm. ASC complex percentage was performed by determining the ratios of cells positives for ASC speckles (Green, GFP) on the total nuclei (Blue, Hoechst). At least 10 fields from n = 3 independent experiments were analyzed. Values are expressed as mean ± SEM. ***p ≤ 0.001, One-Way Anova. F. Plasma membrane permeabilization determination over time using Propidium Iodide (PI) incorporation in WT or NLRP1-deficient pHCECs exposed to Valboro (Vbp, 15µM), EXOA (10ng/mL) or EXOA^H426A^ (10ng/mL) for indicated times. ***p ≤ 0.001, T-test. Values are expressed as mean ± SEM from one experiment (in triplicate) performed at least three times.

EXOA is a ribotoxin that irreversibly ribosylates the elongation factor EEF2, which leads to ribosome inactivation (Jørgensen et al., 2005; Armstrong and Merrill, 2004), a process we also observed by determining ribosome polysomes accumulation (marker of alterations in translation machinery) and translation inhibition (puromycin incorporation as a marker of translation efficiency) (**Fig. 2C, D**). In this context, to determine if EXOA ribosylating activity was required for NLRP1 inflammasome response, we exposed reporter cells to recombinant EXOA and the catalytically dead mutant EXOA^H426A^, which is unable to promote EEF2 ribosylation (Roberts and Merrill, 2002) (**Fig. 2E, S2A**). Microscopy observation and quantifications of NLRP1 inflammasome complexes in reporter cells showed that EXOA but not its mutant triggered robust inflammasome formation (**Fig. 2E**). EEF2 ribosylation by EXOA requires the presence of a specific modified form of Histidine 715 on human EEF2, namely Diphthamide (Liu et al., 2004; Ivankovic et al., 2006) (**Fig. S2B**). This unique amino acid arises from the enzymatic modification of Histidine by the so-called Diphthamide enzymes (DPHs) (Liu et al., 2004; Ivankovic et al., 2006). Hence, using our reporter cells, we genetically deleted *DPH1*, which is critical to initiate diphthamide synthesis (**Fig. S2C**). Analysis of NLRP1 inflammasome complexes in response to EXOA showed that *DPH1*-deficient cells failed to assemble an active NLRP1 inflammasome complex (**Fig. S2D**). As control, activation of the NLRP1 infammasome by the Val-boro-pro (Vbp) molecule was not affected by the removal of DPH1, suggesting that EXOA specifically activates the NLRP1 inflammasome by promoting EEF2-ribosylation on the Diphthamide 715 amino acid (**Fig. S2D**). In addition to EXOA, two other bacterial toxins, namely Diphtheria Toxin (*Corynebacterium diphtheriae*) and Cholix toxin (*Vibrio cholera*) (Jørgensen et al., 2008) also promote irreversible ribosylation of human EEF2 on Diphtamide 715 (Liu et al., 2004; Ivankovic et al., 2006; Jørgensen et al., 2005). Hence, to determine if those toxins could also induce NLRP1 inflammasome formation in a similar way than observed with EXOA, we exposed NLRP1 reporter cell lines to those toxins. As for EXOA, we observed that Cholix Toxin and Diphteria Toxin both triggered robust NLRP1 inflammasome complex assembly in those reporter cells (**Fig. S2D**). Furthermore, NLRP1 inflammasome formation was abrogated in *DPH1*-deficient A549 in response to Cholix Toxin and Diphtheria Toxin, hence confirming the existence of a shared pathway for EEF2 inactivation by those toxins (**Fig. S2D**).

Finally, exposure of primary WT and NLRP1-deficient corneal epithelial cells to EXOA highlighted an increased protection of *NLRP1^-/-^* cells to cell pyroptosis, suggesting that extracellular *P.aeruginosa*-induced NLRP1 inflammasome response in epithelial cells requires T2SS-secreted EXOA and subsequent EEF2 inactivation (**Fig. 2F**).

### EEF2 inactivation drives Ribotoxic Stress Response (RSR)-dependent ZAKα and P38 MAPkinase activation and subsequent NLRP1 inflammasome nucleation

Recent studies unveiled that ribosome inactivation by UV-B as well as the antibiotic anisomycin also promote NLRP1 inflammasome activation in human keratinocytes (Robinson et al., 2022; Jenster et al., 2023; Fenini et al., 2018). Furthermore, this mode of activation engages the MAP3K ZAKα and the effector P38α/β kinases (Robinson et al., 2022; Jenster et al., 2023; Fenini et al., 2018). Thus we wondered if EXOA-driven EEF2a irreversible inactivation and subsequent NLRP1 inflammasome formation might also engages a similar pathway. We generated ZAKα-deficient reporter cell lines and measured their ability to promote activation of the MAPK stress kinases in response to EXOA exposure (**Fig. 3A, S3A**). We observed that EXOA, but not its inactive mutant EXOA^H426A^, strongly induced P38 and JNK stress kinase phosphorylation, a process that disappeared in absence of ZAKα (**Fig. 3A, S3A**). Further analysis of NLRP1 inflammasome formation using fluorescence microscopy highlighted that ZAKα deficiency completely abrogated assembly of the NLRP1 inflammasome and cell death in response to EXOA (**Fig. 3B**). To further analyze if ZAKα and its downstream effectors P38 were important for NLRP1 inflammasome formation, we genetically deleted P38α, P38β or both P38α/β in reporter cells and quantified inflammasome assembly in response to EXOA (**Fig. 3C, S3B**). We observed that single deletion for P38α or P38β did not strongly modify inflammasome assembly whereas combined deficiency for P38α/β led to a robust defect in NLRP1 inflammasome formation upon EXOA exposure (**Fig. 3C**). This suggests that ZAKα- and ZAKα-activated P38 kinases contribute to NLRP1 inflammasome response to EXOA.

**Figure 3.**
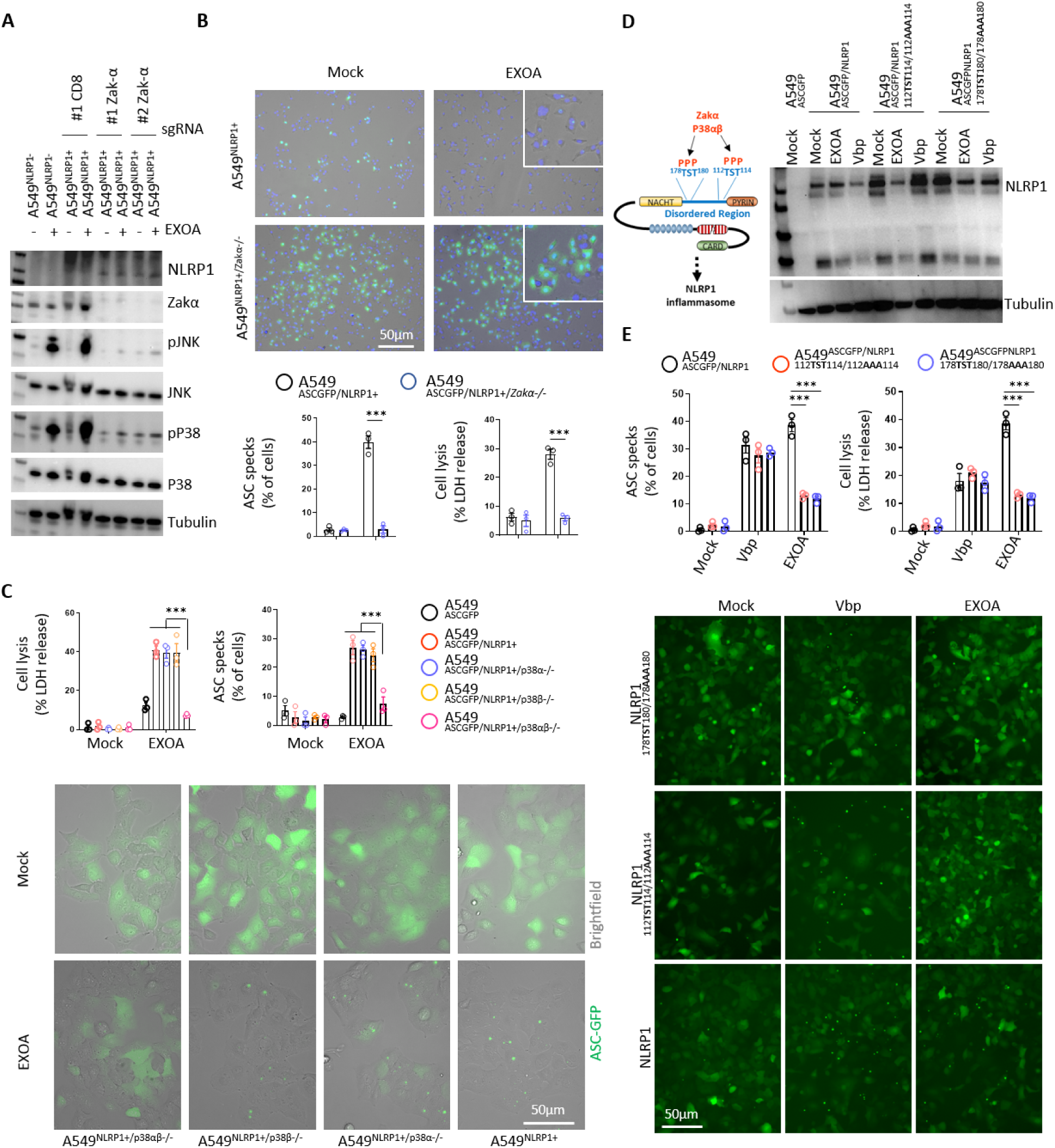
EEF2 inactivation drives ZAKα and P38 MAPkinase activation and subsequent NLRP1 inflammasome nucleation. A. Immunoblotting of P38, JNK, ZAKα, NLRP1, Tubulin and phophorylated P38 and JNK in A549^NLRP1+^ and A549^NLRP1+/ZAKα-^ reporter cell lines exposed or not to EXOA (10ng/mL) for 3 hours. Immunoblots show lysates from one experiment performed at least three times. B. Cell lysis (LDH release), florescence microscopy and associated quantifications of ASC-GFP specks in A549^NLRP1+/ASC-GFP^ and A549^NLRP1+/ASC-GFP/ZAKα-^ reporter cell lines exposed to EXOA (10ng/mL) for 10 hours. ASC-GFP (Green) pictures were taken in dish during after toxin exposure. Images shown are from one experiment and are representative of n = 3 independent experiments; scale bars, 50 µm. ASC complex percentage was performed by determining the ratios of cells positives for ASC speckles (Green, GFP) on the total nucleis (Blue, Hoechst). At least 10 fields from n = 3 independent experiments were analyzed. Values are expressed as mean ± SEM. ***p ≤ 0.001, One-Way Anova. C. Cell lysis (LDH release), florescence microscopy and associated quantifications of ASC-GFP specks in A549^NLRP1+/ASC-GFP^ and A549^NLRP1+/ASC-GFP/P38α/β-^ reporter cell lines exposed to EXOA (10ng/mL) for 10 hours. ASC-GFP (Green) pictures were taken in dish during after toxin exposure. Images shown are from one experiment and are representative of n = 3 independent experiments; scale bars, 50 µm. ASC complex percentage was performed by determining the ratios of cells positives for ASC speckles (Green, GFP) on the total cells (brightfield). At least 10 fields from n = 3 independent experiments were analyzed. Values are expressed as mean ± SEM. ***p ≤ 0.001, One-Way Anova. D. Western blot examination of NLRP1 using an anti-NLRP1 N-terminal antibody (aa 1–323) in A549^ASC-GFP^ reporter cells reconstituted with hNLRP1 or hNLRP1 plasmid constructs mutated for ^112^TST^114^/^112^AAA^114^ or ^178^TST^180^/^178^AAA^180^ after 10 hours exposure to EXOA (10ng/mL) or Vbp (15µM). E. Cell lysis (LDH release), florescence microscopy and associated quantifications of ASC-GFP specks in A549^ASC-GFP^ reporter cells reconstituted with hNLRP1 or hNLRP1 plasmid constructs mutated for ^112^TST^114^/^112^AAA^114^ or ^178^TST^180^/^178^AAA^180^ after 10 hours exposure to EXOA (10ng/mL) or Vbp (15µM). ASC-GFP (Green) pictures were taken in dish during after toxin exposure. Images shown are from one experiment and are representative of n = 3 independent experiments; scale bars, 50 µm. ASC complex percentage was performed by determining the ratios of cells positives for ASC speckles (Green, GFP) on the total cells (brightfield). At least 10 fields from n = 3 independent experiments were analyzed. Values are expressed as mean ± SEM. ***p ≤ 0.001, Two-Way Anova with multiple comparisons.

ZAKα and P38-driven phosphorylation of Serine and Threonines in the NLRP1 disordered region has been recently shown to promote NLRP1 inflammasome activation upon UV-B-driven ribosome collision (Robinson et al., 2022). Here, using NLRP1 constructs mutated for either phosphorylation site 1 (^110^TST^112^) or site 2 (^178^TST^180^), we observed that EXOA was unable to trigger assembly of the NLRP1 inflammasome in reporter cells (**Fig. 3D, E**). To the contrary, Vbp- and SARS-CoV-2-induced NLRP1 inflammasome assembly was fully efficient in cells complemented with either construct or in ZAKα-deficient cells (**Fig. 3D, E, S3C**). All in one, those results suggest that ZAKα- and P38-targeted NLRP1 site 1 and 2 is required for efficient inflammasome assembly in response to EXOA but not during Vbp or SARS-CoV-2 exposure.

### Cystic fibrosis airway epithelial cells show exacerbated sensitivity to EXOA-driven cell death, which is reversed by ZAKα inhibition

Next we investigated the role of EXOA-driven pyroptosis in a patho-physiological model of *P. aeruginosa* infection. In addition to causing corneal infections, *P. aeruginosa* is well known to establish life-threatening infection in airways and lungs of Cystic Fibrosis (CF) patients. Due to defective mucus production and clearance (Veit et al., 2016), the airway of CF patients favor chronic *P. aeruginosa* infections. Collaborating with the Hospital of Toulouse, we obtained nasal brushes from healthy and Cystic fibrosis patients carrying the ΔF508 mutation in the CFTR (ΔF508 / 4005+1G>A and ΔF508 / N1303K, respectively referred in the text and figure legends as CF donors 1 and 2) (**Fig. S4A**). CF patients have been described to express very high basal and inducible P38 kinase activation (Raia et al., 2005; Bérubé et al., 2010), a phenotype we could confirm by exposing or not those cells to EXOA (**Fig. 4A**). Given the importance of P38 kinases at promoting EXOA-dependent NLRP1 inflammasome response, we next measured the ability of healthy and CF nasal cells to undergo EXOA-dependent cell death. Propidium incorporation measure in healthy and CF nasal cells showed that EXOA-driven cell death was exacerbated in CF nasal cells, a process that was reduced by the use of P38 inhibitor SB203580 (**Fig. 4B**). Further experiments also showed that ZAKα inhibition (PLX4720) even further inhibited EXOA-driven cell death, Gasdermin D cleavage and IL-18 release both in healthy and CF nasal cells (**Fig. 4C-F, S4B**). Such process was conserved with the ribotoxic antibiotic anisomycin but not in response to Vbp where nasal cells from healthy and CF patients responded similarly, hence suggesting that P38 pre-activation in CF patients specifically sensitive them to ribotoxic stress-driven NLRP1 response, a process that may contribute to *P. aeruginosa*-driven epithelial barrier disruption (**Fig. 4D-F, S4B**). Altogether, our results suggest that the ZAKα/NLRP1 axis contributes to exacerbated epithelial barrier destabilization upon EXOA-inactivated EEF2 exposure.

**Figure 4.**
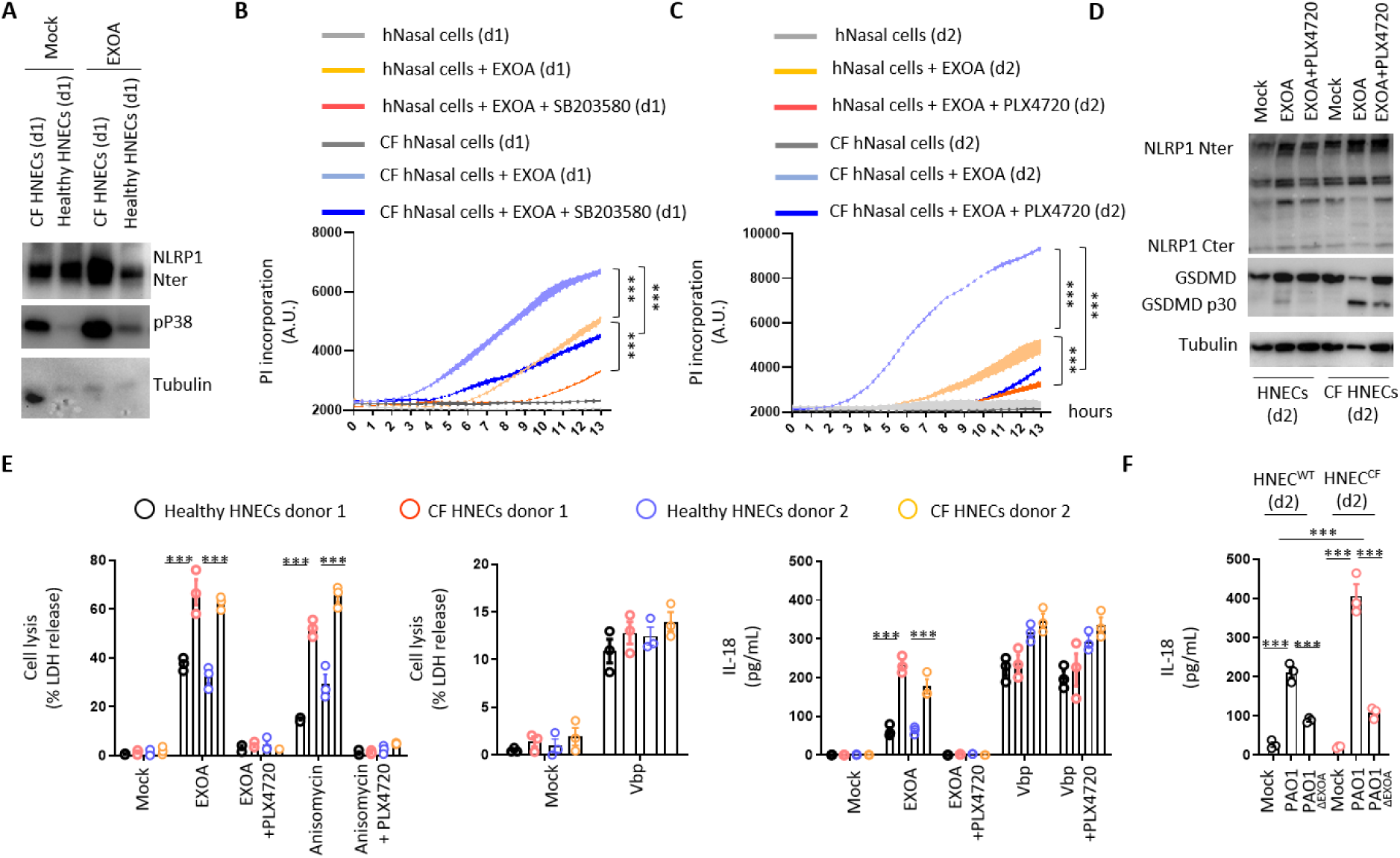
Cystic fibrosis airway epithelial cells show exacerbated sensitivity to EXOA-driven pyroptosis, which is reversed by ZAKα inhibition. A. Immunoblotting of NLRP1, Tubulin and phophorylated P38 in pHNECs^WT^ and pHNECs^CF^ from healthy (WT) and Cystic Fibrosis (CF) patients exposed to EXOA (10ng/mL) or not for 12 hours. Immunoblots show lysates from one experiment performed at least three times. (d1) stands for “donor 1” from Cystic fribosis (CF) or healthy (WT) patients. B, C. Plasma membrane permeabilization determination over time using Propidium Iodide (PI) incorporation in pHNECs^WT^ or pHNECs^CF^ exposed to EXOA (10ng/mL) for indicated times. When specified, SB203580, inhibitor of P38 activity (10µM) or PLX420 (bRaf, ZAKα inhibitor, 10µM) were used. (d1) and (d2) stand for respective donors 1 or 2. ***p ≤ 0.001, T-test. Values are expressed as mean ± SEM from one experiment (in triplicate) from one independent donor (d1/d2, CFd1/CFd2) performed at least three times. D. Immunoblotting of NLRP1, Gasdermin D (GSDMD) and Tubulin in pHNECs^WT^ and pHNECs^CF^ from healthy (WT) and Cystic Fibrosis (CF) patients exposed to EXOA (10ng/mL) or not for 12 hours in presence or absence of PLX420 (ZAKα inhibitor, 10µM). Immunoblots show combined supernatants and lysates from one experiment performed at least three times. (d2) stands for “donor 2” from Cystic fribosis (CF) or healthy (WT) patients. E. Cell lysis (LDH) and IL-18 release evaluation in pHNECs^WT^ and pHNECs^CF^ upon EXOA (10ng/mL), Anisomycin (1µg/mL) or Vbp (15µM) treatement for 18 hours in presence/absence of PLX420 (ZAK inhibitor, 10µM). ***p ≤ 0.001, T-test. Values are expressed as mean ± SEM from one experiment (in triplicate) from one independent donor (d1/d2, CFd1/CFd2) performed at least three times. F. IL-18 release in pHNECs^WT^ and pHNECs^CF^ co-cultured with PAO1 or PAO1^ΔEXOA^ (1.10^5^ bacteria) for 24 hours. ***p ≤ 0.001, T-test. Values are expressed as mean ± SEM from one experiment (in triplicate) from one independent donor (d2, CFd2) performed at least three times.

## DISCUSSION

Chronic infections of multiple organs mediated by *P. aeruginosa* are a major source of epithelium damage and inflammatory exacerbations. Due to the lack of robust and relevant models of study as well as its remarkable adaptation to specific environments, the impact of chronic infections mediated by *P. aeruginosa* on host epithelial integrity has long been hard to address. Here, our findings and those from the Zhong group (companion manuscript) that the T2SS-released Exotoxin A, and its relatives Cholix and Diphtheria Toxins, trigger activation of the human NLRP1 inflammasome in skin keratinocytes, corneal and airway epithelial cells, three important sites of *P. aeruginosa* chronic infections. This process exemplifies a novel biochemical pathway by which epithelial organs detect bacterial virulence factors.

Although major research studies unveiled that upon acute infection, *P. aeruginosa* T3SS allows both activation of the NLRC4 and NLRP3 inflammasomes in rodent as well as in human macrophage and neutrophil models (Sutterwala et al., 2007; Faure et al., 2014; Franchi et al., 2007; Miao et al., 2008; Deng et al., 2015; Balakrishnan et al., 2018; Santoni et al., 2022a; Ryu et al., 2016), chronic infections mediated by this pathogen is associated with a downregulation of T3SS in favor a biofilm phenotype, where EXOA is strongly produced and released. To this regard, our observation that EXOA-driven ribotoxic stress contributes to exacerbated tissue damage and inflammation strongly correlate with earlier studies which showed that EXOA-deficient bacteria triggered lower tissue damages during infections of human and mice (Michalska and Wolf, 2015; Pillar and Hobden, 2002). Even though the RSR-driven NLRP1 inflammasome path is not conserved among rodents and humans, ZAKα-driven RSR constitutes a shared process between both species, which suggests that evolution selected ZAKα as a versatile stress sensor. To the contrary, for yet to be determined reasons, human-specific evolution has linked RSR to NLRP1 inflammasome response in epithelia, which suggests a specific importance for such selection.

A caveat of this work mostly relies on the yet to be determined identification of Ubiquitin ligases that promote NLRP1 functional degradation upon ZAKα-driven phosphorylation, which is currently under investigations. The sensitivity of NLRP1 activation to the Nedd8 inhibitor MLN4924 suggests that Nedd8-driven Cullin ligase activation is of major importance in this process (Jenster et al., 2023; Robinson et al., 2022). Among the broad family of Cullin ligases, Cull1 and 2 were found to target phosphorylated proteins for Ubiquitination and subsequent degradation (Chen et al., 2021). Should one of those Ubiquitin ligase complex be involved constitutes an attractive hypothesis to pursue.

In addition to UVB, Chikungunya virus, the antibiotic anisomycin and the fungal toxin DON (Fenini et al., 2018; Jenster et al., 2023; Robinson et al., 2022), our work, along the one from Zhong lab, unveils the critical involvement of the NLRP1 inflammasome upon infections mediated by various bacterial pathogens, including *C. diphtheria* and *P. aeruginosa*. To this regard, patients developing Cystic fibrosis (CF) show exacerbated inflammation and tissues damages upon chronic infection with *P. aeruginosa*. Specific studies highlighted that stress-activated kinases P38 and JNK were over activated in CF-derived cells from patients (Bérubé et al., 2010; Raia et al., 2005). Although various hypothesis were developed to explain such dysregulation in CF patients, no study, including ours, could unveil the critical molecular and biochemical mechanisms engaged, which warrants for further investigations. However, our findings that in this context, EXOA-but also other RSR inducers specifically triggered an exacerbated cell death response suggest that targeting ZAKα and/or P38 kinases in *P. aeruginosa*-infected patients might constitute a good host-targeted approach in order to limit epithelial damages complementary to the current antibiotic and CFTR modulator strategy used (Kaftrio | European Medicines Agency).

Finally, it is long been noted that host EEF2 (and EEF1) is targeted by a variety of exotoxins from different unrelated bacterial species. Indeed, EEF2 is highly conserved in all eukaryotic species, all expressing a specific diphtamide amino acid. In this context, yeasts as well as mammals are similarly targeted by EXOA and relative toxins, which underlines the outstanding adaptation of *P. aeruginosa* to its environment but also raises the question of the specific species where EXOA holds the most prevalent/potent role for bacterial development/survival. Another key question lies on the recent identification DPH1 and 2 deficient patients (Urreizti et al., 2020; Nakajima et al., 2018; Hawer et al., 2020). If those deficiencies might actually confer a selective advantage or not to *C. diphteria*, *V. cholera* or *P. aeruginosa* infections will constitute an exciting field of investigations, as previously observed for HIV- or-malaria-resisting patients (Samson et al., 1996; Allison, 1954) or recently highlighted for *Yersinia pestis*-shaped selection/evolution of the inflammasome-forming sensor PYRIN (Park et al., 2020).

All in one, our results describe the critical role of ZAKα-driven NLRP1 inflammasome response and epithelial disruption in response to the pathogen *P. aeruginosa*, and exemplify its deleterious potential in CF pathogenesis.

## ACKNOWLEDGMENTS

We are extremely grateful to patients and their families for their willingness of participating to this study. Authors also warmly acknowledge Y. Rombouts, L. Boyer, T. Henry, M. Tiraby and all lab members for their outstanding advices and support on this project.

This project was supported by the ATIP-Avenir program (to EM), FRM “Amorçage Jeunes Equipes” (AJE20151034460 to EM), the Agence Nationale de la Recherche (ANR Psicopak to EM), the ANRS-MIE (to EM), the ERC (StG INFLAME 804249 to EM), the European Society of Clinical Microbiology and Infectious Diseases (ESCMID, to RP), Invivogen-CIFRE PhD grant (to MP), Vaincre La Mucoviscidose (VLM) and Region Occitanie (GRAINE) grants to CC.

The funders had no role in study design, data collection and analysis, decision to publish, or preparation of the manuscript.

## MATERIAL AND METHODS

### Ethic statements

Patient data and tissue collection was performed in agreement with European Network of Research Ethics Committees and French ethic law. The ethical committee, according to the Medical Research Involving Human Subjects Act, reviewed and approved the study. Human tissue was provided by the University Hospital of Toulouse (France) and CNRS (agreements CHU 19 244 C and CNRS 205782). All patients involved in this study declared to consent to scientific use of the material; patients can withdraw their consent at any time, leading to the prompt disposal of their tissue and any derived material.

### Reagent used in the study

| Antibody | Catalog Reference | Provider |
| --- | --- | --- |
| Rabbit anti-ZAK antibody 1: 1000 | A301-993A | Bethyl Laboratories |
| P38 MAPK antibody 1: 1000 | 9212S | Cell signaling |
| Phospho-P38 MAPK (Thr180/Tyr182) (D3F9) XPRabbit mAb 1: 1000 | 4511S | Cell signaling |
| JNK1 Rabbit anti-Human Polyclonal 1: 1000 | 44690G | Invitrogen |
| Phospho-SAPK/JNK (Thr183/Tyr185) (81E11) Rabbit mAb 1: 1000 | 4668S | Cell signaling |
| Anti NLRP1 1:200 | 679802 | Biolegend |
| Anti NLRP1 (N-terminal) 1: 1000 | AF6788-SP;RRID: AB_2916167 | R&D systems |
| Anti- NLRP1 (C-terminal), 1: 1000 | ab36852;RRID: AB_776633 | abcam |
| anti-Sheep igG HRP (1/4000) | HAF016;RRID: AB_562591 | R&D |
| Goat-anti-rabbit IgG (H+L), HRP conjugate (1/5000) | R-05072-500;RRID: AB_10719218 | Advansta |
| GSDMDC1 antibody 1: 1000 | NBP2-33422 | biotechne |
| Anti-Puromycin Antibody, clone 12D10 | MABE343 | Sigma |
| EEF2 antibody (1:1000) | 2332S | Cell signaling |
| DPH1 antibody (1:1000) | # H00001801-M02 | ThermoFisher scientific |
| Anti-alpha Tubulin antibody 1: 1000 | ab4074 | Abcam |
| Streptavidin - HRP | 434323 | ThermoFisher scientific |
| **Reagents** | **Catalog Reference** | **Provider** |
| Transwell | 140620 | ThermoFisher scientific |
| NAD<sup>+</sup>, Biotin-Labeled | 80610 | BPSBioscience |
| Pseudomonas exotoxin A 0.5 mg | P0184-.5MG | sigma |
| Inactive Exotoxin A | N.A. | Creative Diagnostics |
| Diphtheria Toxin | D0564-1MG | Sigma |
| Cholix toxin | This study | This study |
| Anisomycin | SE-S7409-10MG | selleck |
| Talabostat (Vbp) | tlrl-vbp-10 | invivoGen |
| PhoSTOP | 4906845001 | sigma/Roche |
| cOmplete, Mini, EDTA-free Protease Inhibitor Cocktail | 4693159001 | Sigma/roche |
| Lipofectamine LTX | 15338030 | Invitrogen |
| Puromycin Dihydrochloride | A1113803 | gibco |
| Molecular Probes SYTOX Green Nucleic Acid Stain | S7020 | Thermofisher |
| Propidium iodide | P1304MP | Thermofisher |
| Nate | lyec-nate | Invivogen |
| **Critical commercial assays** | **Catalog Reference** | **Provider** |
| Human total IL-18 duo set | DY318-05 | R&D |
| Cyquant LDH | C20301 | Thermofisher |
| Human IL-1beta | 88-7261-77 | Thermofisher |
| Q5 Site-Directed Mutagenesis Kit Protocol | E0554 | NEB |
| **Chemicals** | **Catalog Reference** | **Provider** |
| ZAK inhibitor (PLX-4720) 10µM | HY-51424 | Med Chem Express |
| P38 inhibitor SB 203580 10µM | HY-10256 | Med Chem Express |
| Caspase-1 inhibitor - Ac-YVAD-cmk 20µM | inh-yvad | invivogen |
| Z-VAD-FMK 20µM | vad-tlrl: | invivogen |
| Caspase-3/7 inhibitor ZDEVD-FMK 20µM | S7312 | Selleck |
| Caspase-8 inhibitor - Z-IETD-FMK | inh-ietd | invivogen |
| Bortezomib 1µM | S1013 | selleck |
| Cycloheximide (CHX) | C4859-1ML | Sigma |
| **Human Primary Nasal Epithelial cells** | **Catalog Reference** | **Provider** |
| Tryple Express Enzyme (1X), no phenol red | 10718463 | Fisher Scientific |
| Nutragen Type I Collagen Solution, 6 mg/ml (Bovine) | 5010-50ML | Advanced Biomatrix |
| Sputolysin | 560000-1SET | Sigma |
| PneumaCult-Ex Plus Medium | 05040 | Stem cell |
| Costar 6.5 mm Transwell, 0.4 µm Pore Polyester Membrane Inserts | 38024 | Stem cell |
| **Human Primary Corneal epithelial cells** | **Catalog Reference** | **Provider** |
| Human Corneal Epithelial Cell Growth Medium | 221-500 | Tebu-bio |
| Human corneal epithelial cells | 630-05a | Tebu-bio |
| **Cell lines** | **Catalog Reference** | **Provider** |
| A549 ASC GFP NLRP1 | a549-ascg-nlrp1 | Invivogen |
| A549 ASC GFP | a549-ascg | Invivogen |
| A549 ASC GFP NLRP1/ACE2 | a549-ascov2-nlrp1 | Invivogen |
| A549 ASC GFP ACE2 | a549-ascov2 | Invivogen |
| **Recombinant DNA** | **Catalog Reference** | **Provider** |
| hNLRP1 gene | puno1-hnalp1a | Invivogen |

### Cell culture

A549 cells were maintained in Dulbecco’s modified Eagle’s medium (DMEM,Gibco) supplemented with 10% heat-inactivated fetal bovine serum (FBS), 1% penicillin-streptomycin, and 1% L-glutamine at 37°C 5% CO2. Primary Human Corneal epithelial cells were maintained in Human Corneal Epithelial Cell Growth Medium (Tebu-bio) at 37°C 5% CO2.

Regarding primary Human Nasal Epithelial Cells (pHNECs), patients’ pHNECs were collected on superior turbinates using smear brushes at the Hosipital of Toulouse, France. After brushing back cells in collection medium, centrifugation was performed for 5 min 400g at 4°C. Pellet was resuspended in 4 ml TrypLE express (GIBCO) + 20µl Sputolysin (200X) and incubated at 37°C for 5 minutes to disrupt mucus. TrypLE was diluted with 4 ml of Advanced DMEM F12-.

Pellet was recovered after centrifugation and culture was continued in the expansion medium Pneumacult). After a week of proliferation, basal cells were counted and seeded onto collagen-coated (0.03mg/mL) and maintained in Pneumacult Ex Plus Medium (StemCell) at 37°C 5% CO2.

### Cell Stimulations

Otherwise specified, cells were plated one day before stimulation in 12-well plates at 200 000 cells per well in 1mL of DMEM, 10% FCS, 1% PS.

Medium was changed to OPTIMEM and cells were preincubated or not with the indicated inhibitors during 1 hour.

The pHNE cells were seeded the day before the stimulation in 12-well plates at 200 000 cells per well in 1mL of pneumacult medium. Cell’s medium was changed to OPTIMEM and cells were treated or not with the indicated inhibitors during 1 hour.

The pHCE cells were seeded the day before the stimulation in 12-well plates at 10 000 cells per well in 1 mL of corneal epithelial cell growth medium. Medium of the cells was changed to OPTIMEM and cells were treated or not with the indicated inhibitors during 1 hour.

All cells were treated with the indicated concentration of Exotoxin A (EXOA, 10 ng/mL to 1µg/mL), anisomycin (1µM) or with Val-boroPro (15µM) for indicated times.

SARS-CoV-2 (BetaCoV/France/IDF0372/2020 isolate) experiments were performed in BSL-3 environment as described in (Planès et al., 2022).

Briefly, 250,000 A549^ASC-GFP/NLRP1/ACE2^, genetically invalidated or not for ZAKα, were infected for 24 hours with BetaCoV/France/IDF0372/2020 strain at indicated MOI in DMEM supplemented with 10mM Hepes, 1% penicillin-streptomycin and 1% L-Glutamine for 1h at 37°C.

### Transwell infections

A549 cells, expressing or not NLRP1, along with the ASC-GFP reporter, were plated in 24 well plates at 2, 5.10^5^ cells per well one day before experiment. The following day, cell culture media was replaced by 600 µL of OPTI MEM per wells. Cells were placed in co-culture with Pao1 bacteria WT or mutant (as indicated) at MOI of 10, in 100 µL of OPTI MEM separated by a semi-permeable transwell insert (0,3µm). After 18 hrs of co-culture, transwell inserts containing bacteria were removed and ASC-speck formation in A549 cell was analyzed. Images were acquired using EVOS M700 microscope.

### Bacterial growth and mutants

*P. aeruginosa* strains (PAO1) and their isogenic mutants were grown in Luria Broth (LB) medium overnight at 37°C with constant agitation. The following day, bacteria were sub-cultured by diluting overnight culture 1:25 and grew until reaching an optical density (OD) O.D.600 of 0.6 – 0.8.

PAO1 and its transposon mutants were obtained from two-allele transposon library (Jacobs et al., 2003).

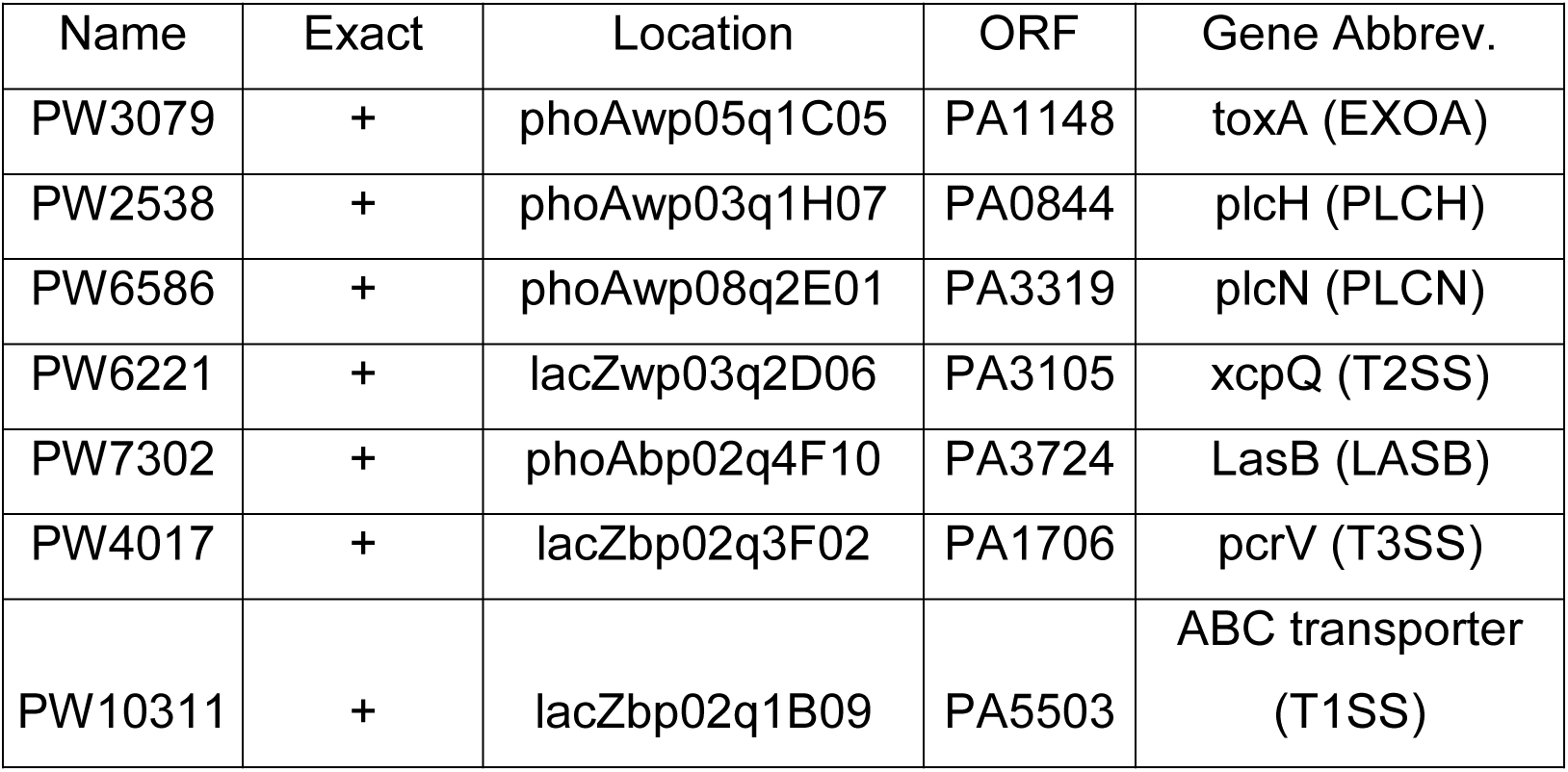

| Name | Exact | Location | ORF | Gene Abbrev. |
| --- | --- | --- | --- | --- |
| PW3079 | + | phoAwp05q1C05 | PA1148 | toxA (EXOA) |
| PW2538 | + | phoAwp03q1H07 | PA0844 | plcH (PLCH) |
| PW6586 | + | phoAwp08q2E01 | PA3319 | plcN (PLCN) |
| PW6221 | + | lacZwp03q2D06 | PA3105 | xcpQ (T2SS) |
| PW7302 | + | phoAbp02q4F10 | PA3724 | LasB (LASB) |
| PW4017 | + | lacZbp02q3F02 | PA1706 | pcrV (T3SS) |
| PW10311 | + | lacZbp02q1B09 | PA5503 | ABC transporter<br>(T1SS) |

Specific deletion of xcpQ (T2SS) and toxA (EXOA) genes was achieved as described previously in (Santoni et al., 2022b). Briefly, pEXG2 suicide vector containing 700-bp sequences of the flanking regions of the selected gene was directly inserted into competent SM10λpir (Mix&Go competent cells, Zymo Research Corporation) and subsequently selected on LB-Agar supplemented with 50 μg/mL kanamycin /15 μg/mL gentamicin. After sequencing resulting clones were mated with PAO1 strains, 4H/37°C. Mated bacteria were plated on 15 μg/mL gentamicin and 20 μg/mL Irgasan LB-Agar plates in order to selectively remove E.coli SM10 strains. Next day, 5-10 clones were grown for 4h in LB and streaked on 5% sucrose LB plates overnight at 30°C. PAO1 clones were then checked by PCR for mutations.

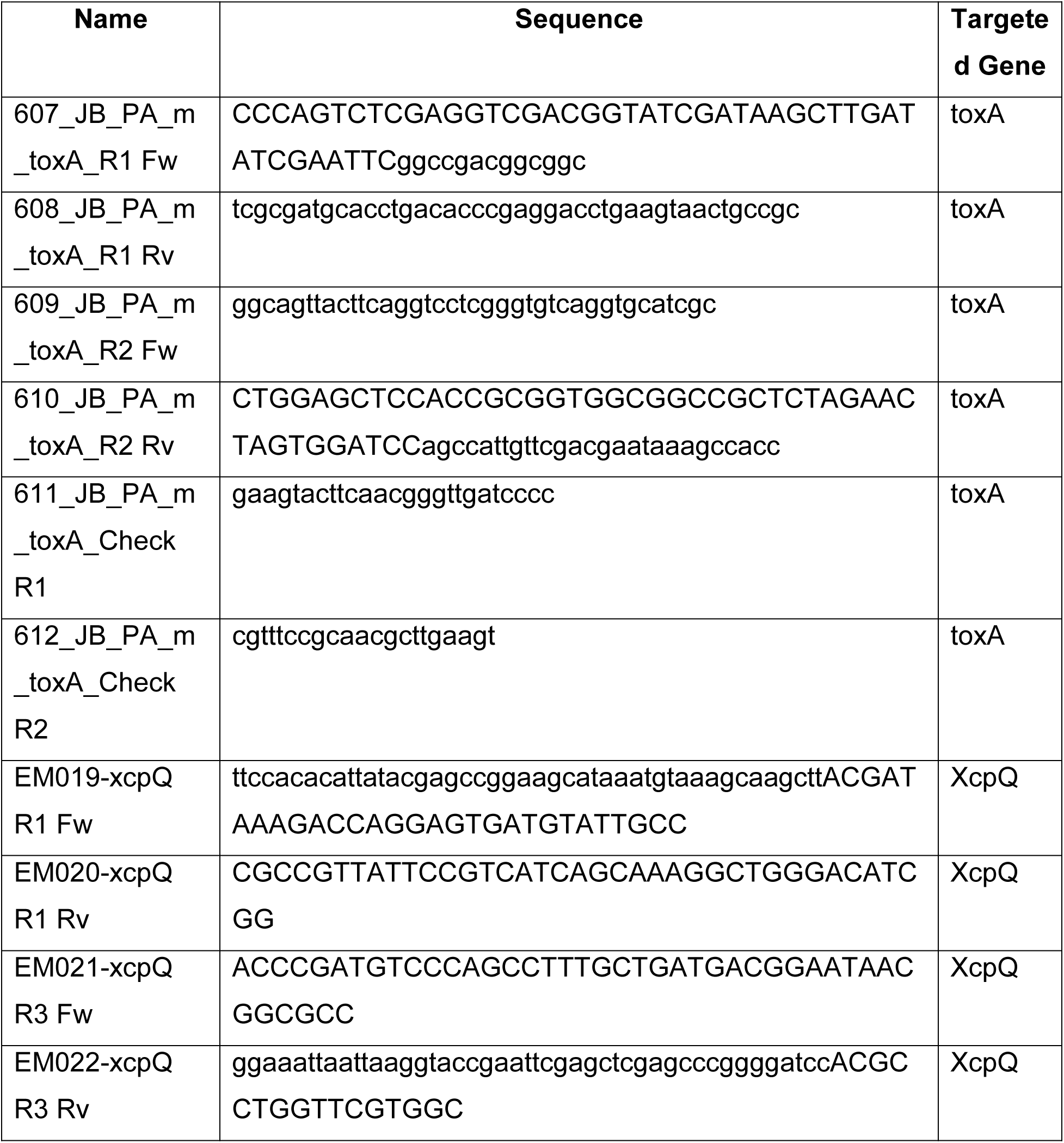
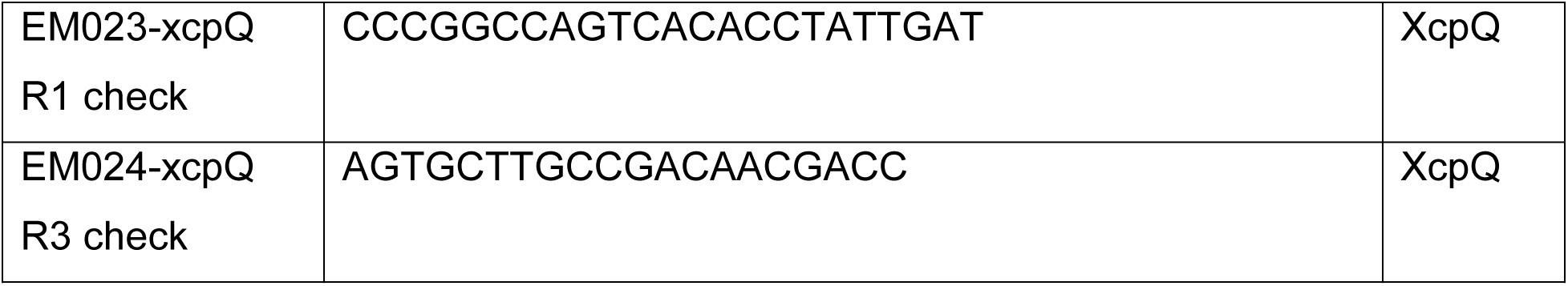

### Cell death assays

Cell lysis was measured by quantification of the lactate dehydrogenase (LDH) release into the cell supernatant using LDH CyQUANT kit (Thermofisher). Briefly, 50 µL cell’s supernatants were incubated with 50 µL LDH substrate and incubated for 30 min at room temperature protected from light.. The enzymatic reaction was stopped by adding 50 µL of stop solution. Maximal cell death was determined with whole cell lysates from unstimulated cells incubated with 1% Triton X-100

### Plasma membrane permeabilization assays were performed using propidium iodide incorporation

Cells were plated at density of 1 x 10^5^ per well in Black/Clear 96-well Plates in OPTI-MEM culture medium supplemented with Propidium Iodide dye (1µg/mL) and infected/treated as mentioned in figure legends. Red fluorescence is measured in real-time using a Clariostar plate reader equipped with a 37°C cell incubator or using an EVOS Floid microscope (Invitrogen). Maximal cell death was determined with whole cell lysates from unstimulated cells incubated with 1% Triton X-100.

### Cytokine quantification

Cytokines secretions was quantified by ELISA kits, according to the manufacturer’s instructions, IL-1B (Thermo Fisher Scientific, (88-7261-77), IL-18 (Biotechne, DY318-05).

### Sample preparation for immunoblot

At the end of the experiment, cell’ supernatant was collected and soluble proteins were precipitated using trichloroacetic acid (TCA) as described previously (Santoni et al., 2022b). Precipitated pellet was then resuspended in 50 uL of RIPA buffer (150 mM NaCl, 50 mM Tris-HCl, 1% Triton X-100, 0.5% Na-deoxycholate) supplemented with protease inhibitor cocktail (Roche). Adherent cells were lysed in 50 uL of RIPA buffer supplemented with protease inhibitor cocktail (Roche). Cell lysate and cell supernatant were homogenized by pipetting up and down ten times and supplemented with laemli buffer (1X final) before boiling sample for 10 min at 95°C.

### Immunoblot

Cell lysates were separated by denaturing SDS-PAGE and transferred on PVDF membrane. After transfer, the membrane is saturated 1h at room temperature in TBS-T (Tris 10 mM pH 8, NaCl 150 mM, Tween 20 0.05%) containing 5% BSA. Then, the membrane is incubated overnight at 4°C with the different primary antibodies, under agitation. After 3 washes with TBS-T, the membrane is further incubated with the secondary antibodies coupled with the peroxidase enzyme HRP (horseradish peroxidase) for 1 hour at room temperature and under agitation. Then membranes are washed 3 times with TBS-T. ECL revelation kit (Advansta) was used as a substrate to reveal HRP activity and membranes are imaged using ChemiDoc Imaging System (BioRad). The primary antibodies and secondary antibody used are listed in Reagent table.

### Phosphoblots

At the end of the experiment, cell’ supernatant was discarded and adherent cells were lysed in 50 uL of RIPA buffer (150 mM NaCl, 50 mM Tris-HCl, 1% Triton X-100, 0.5% Na-deoxycholate) supplemented with protease inhibitor cocktail (Roche) and phosphatase inhibitors cocktails (Roche). Collected cell lysate was homogenized by pipetting up and down ten times and supplemented with laemli buffer before boiling for 10 min at 95°C. Cell lysates were then separated by SDS-PAGE and handled as described in the immunoblot section.

### Puromycin incorporation

The day before the stimulation A549 ASC-GFP expressing or not NLRP1 were seeded in 12-well plates at 200 000 cells in 1mL of DMEM 10% FCS 1% PS. Cells were treated with exotoxin A (10 ng/mL) for indicated times. 30 min before the end of the treatment puromycin antibiotic was added in cell medium at 1ug/mL final. Following puromycin incubation, supernatant was discarded and adherent cells were prepared for immunoblot as described in the immunoblot section. Puromycin incorporation was revealed using the following antibody Anti-Puromycin Antibody (clone 12D10 MABE343).

### Polysomes Profiling

Sucrose gradient preparation. Five sucrose solutions containing 10 %, 20 %, 30 %, 40 % and 50 % sucrose (w/v) were prepared in TMK buffer (20 mM Tris-HCl pH 7.4, 10 mM MgCl2, 50 mM KCl). Layers of 2.1 mL of each solution were successively poured into 12.5 mL polyallomer tubes (Beckman-Coulter, Cat. # 331372), starting from the most concentrated solution (50 %) at the bottom of the tubes to the least concentrated solution (10 %) at the top. Each layer was frozen in liquid nitrogen before pouring the following one. Frozen gradients were stored at - 80 °C and slowly thawed overnight at 4°C before use. Extract preparation. A549 cells were grown to 70% confluency and then treated or not with exotoxin A (10 ng/mL) for indicated times. Following this treatment, cells were incubated with Cicloheximide (CHX, 100 ug/mL) for 15 min at 37°C. Following treatment, cells were rinsed twice with PBS and treated with 1mL of trypsin (0,25%) for 5 min at 37°C. Trypsin was diluted with DMEM medium containing 100 ug/mL of CHX. Cells were mixed up and down and counted to adjust the final resuspension volume in each condition. Cells were centrifuged at 1200 rpm (300g) for 5 min at 4°C. The cell pellet was washed with ice-cold PBS containing 100ug/mL of CHX. Cells were centrifuged at 1200 rpm (300g) for 5 min at 4°C. Supernatants were aspirated and gently lysed in lysis buffer (20 mM Tris-Cl [pH 8], 150 mM KCl, 15 mM MgCl2, 1% Triton X-100, 1mM DTT, 100ug/mL CHX, EDTA-free and Protease inhibitor cocktail. Samples were incubated on ice for 20 min and centrifuge at 1000 g for 5 min at 4 °C min and supernatant corresponding to the cytosolic fraction of the cells were collected into a 1.5 mL tube. Samples were further centrifugated at 10 000 g for 5 min at 4°C to clarify the cytoplasmic extract. Extracts were quantified by measuring absorbance at 260 nm using Nanodrop.

### Extracts loading, gradient centrifugation and collection

Normalized amounts of extracts were loaded on 10–50% sucrose gradients, and then centrifuged at 260,800 x g for 2.5 h at 4°C in an Optima L-100XP ultracentrifuge (Beckman–Coulter) using the SW41Ti rotor with brake. Following centrifugation, fractions were collected using a Foxy R1 gradient collector (Teledyne Isco) driven by PeakTrak software (Version 1.10, Isco Inc.). The A_254_ was measured during collection with a UA-6 UV/VIS DETECTOR (Teledyne Isco). The final polysome profiles were generated in Excel from .txt files extracted from PeakTrak software.

### EEF2 ribosylation assays

Cell lysates from A549 were prepared by suspending one pellet of 1.10^6^ A549 cells in 100 µL of RIPA buffer. For in-vitro ADP-ribosylation assays, reactions were performed in Eppendorf tube by mixing 50 µL of whole cell lysate with 100 µL of ADP-ribosylation buffer [20 mM Tris-HCl (pH 7.4), 150 mM NaCl and 1 mM DTT] supplemented with NAD+ Biotin-Labeled (BPSBioscience) at 50 µM final, in presence or absence of recombinant EXOA protein (100 ng). Reaction was left for 1h at 25°C. 20 µL of the reaction was analyzed by SDS-PAGE followed by Western blotting as described in the immunoblot section. Membranes were first subjected to EEF2 detection (using anti-EEF2 followed by HRP-conjugated secondary antibody), then membranes were stripped, and ADP-ribosylation was detected by monitoring the incorporation of the NAD+ Biotin-Labeled probe using streptavidin-HRP-conjugate.

### Generation of mutations in NLRP1 gene

To generate NLRP1 gene mutated for each phosphorylation sites (site 1 (^110^TST^112^) or site 2 (^178^TST^180^)) Threonines and Serines were substituted with Alanine in the human NLRP1 gene (isoform 1) by site-directed mutagenesis using Q5 site-directed mutagenesis Kit Protocol (E0554) according to the manufacturer’s instructions. Following primers were used:

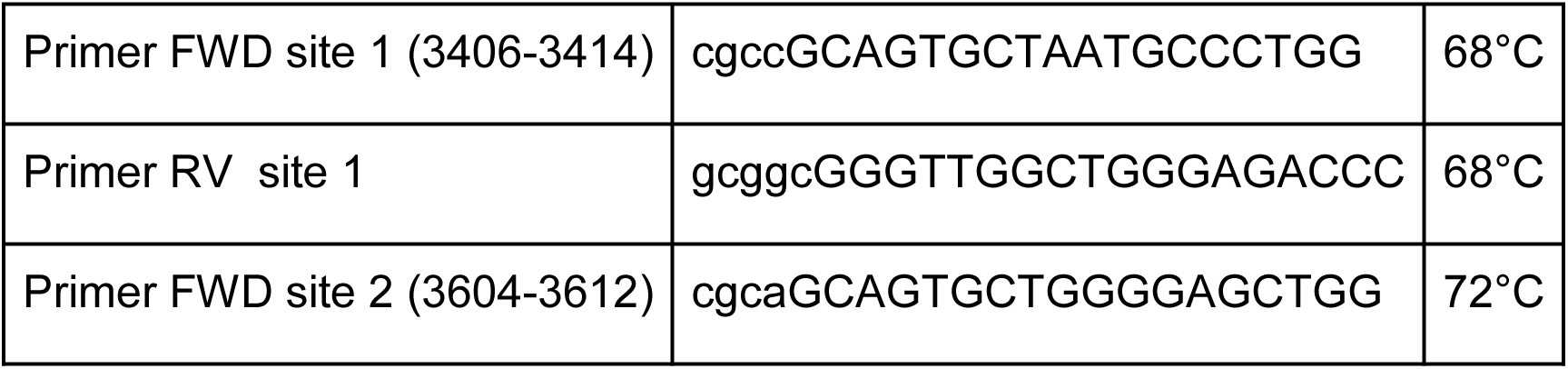
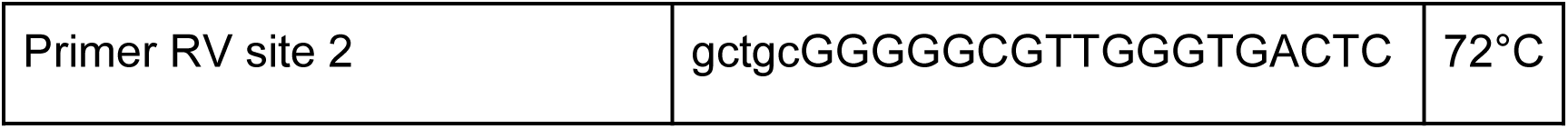

### Cell transfection

The day prior to the transfection, A549 cells were plated in 6 well plate at 2 x 10^5^ cells per well in 1ml of DMEM complete medium. The following day, cells were incubated with Nate 1X (Invivogen) for 30 min. 1 µg of NLRP1 plasmids (pLvB72 hNLRP1, pLvB72 hNLRP1(3604-3612), pLvB72 hNLRP1 (3406-3414) were transfected using lipofectamine LTX and PLUS reagent according to the manufacturer’s instructions (Invitrogen). Transfected cells were incubated for 48 h before further treatments.

### Cholix toxin production

Recombinant cholix toxin (ChxA) was produced using an adapted methodology from (Ogura et al., 2011). Briefly, BL21 (DE3) *E. coli* expressing chxA with a N-terminus hexahistidine-MBP-tag were harvested and bacteria were lysed by sonication on ice. Recombinant cholix toxin was first purified on nickel metal affinity chromatography (Takara) and subsequent TEV protease-mediated 6His-MBP tag removal at 4°C/Over Night (ON). Then, a second nickel metal affinity chromatography allowed harvesting Cholix toxin in the flow-through fraction.

### Genetic invalidations

Genetic invalidation of NLRP1 in primary Human Nasal Epithelial Cells (pHNEs) and primary Human Corneal Epithelial Cells (pHCECs) was achieved by using Ribonucleoprotein (RNP) technic and nucleofection as described previously (Planès et al., 2022). RNP mixes containing Cas9 protein (90pmoles, 1081059, IDT), gRNA (450pmoles) and electroporation enhancer (1μL/Mix, 1075916, IDT) were electroporated using the Neon transfection system (Life Technologies) in T Buffer (Life Technologies). Settings were the following: 1900 V Voltage, 10 Width, 1 Pulse, 20ms.

Efficient NLRP1 targeting sgRNA sequence was provided by FL. Zhong (5′ GATAGCCCGAGTGCATCGG 3′) (Robinson et al., 2022).

Regarding genetic invalidation of P38 isoforms and ZAKα, A549 cells were transduced with LentiCRISPR-V2 vectors containing sgRNA guides against DPH1, P38 isoforms and ZAKα. 48h after transduction, cells were selected during 2 weeks and puromycin double resistant cells were used in functional assays. A second round of lentiviral infection with lenti-Blasticidin-sgP38β in single KO cells for P38α achieved generation of double KO cells for P38α and P38β. Cells were subsequently selected in blasticidin antibiotic before checking genetic invalidation by immunoblotting.

Guides used to generate genetic invalidation

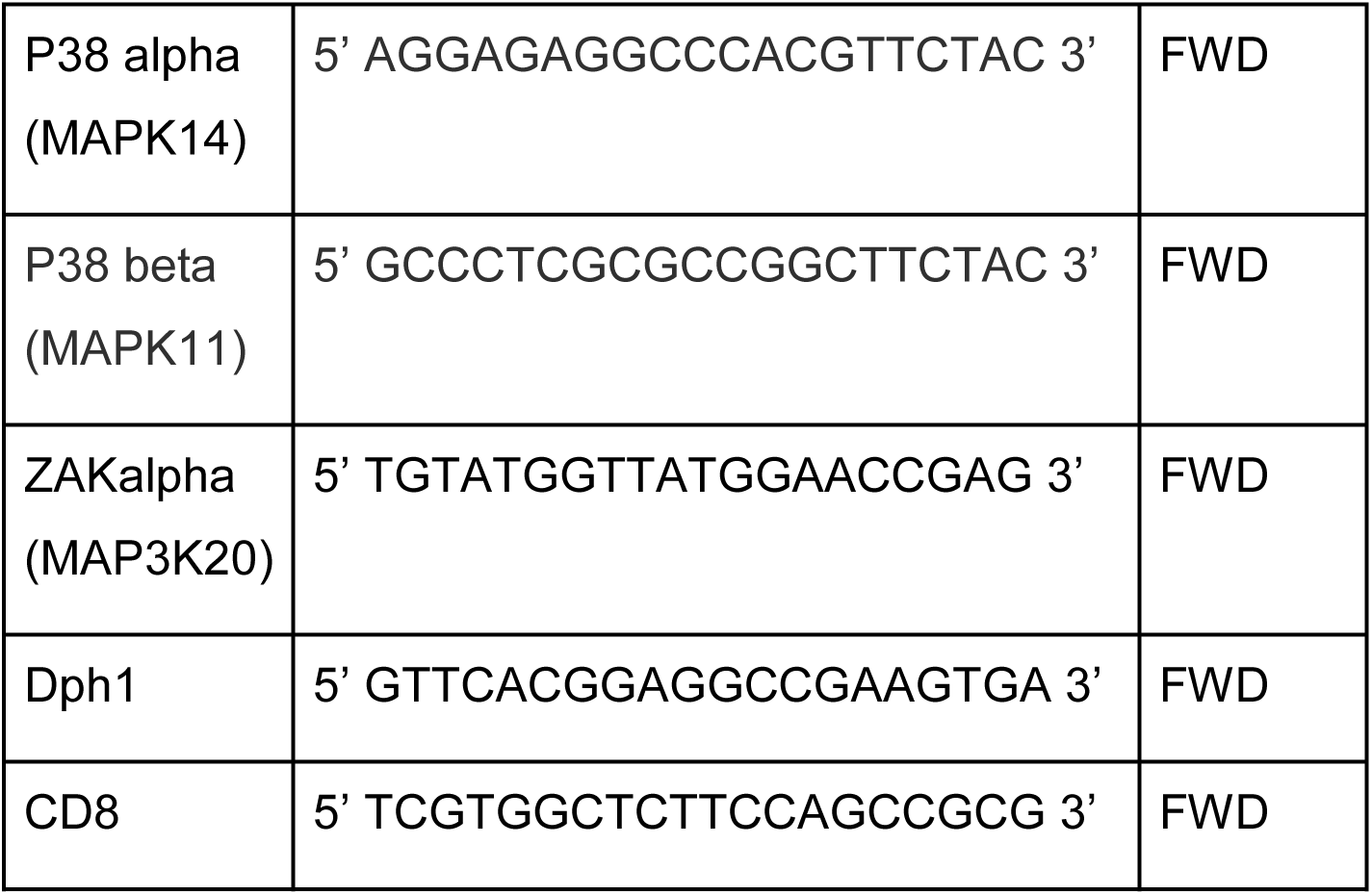

### Analysis

Prism 8.0a (GraphPad Software, Inc) wa used to perform statistical analysis. All revlevant informations are included directly in figure legends. Otherwise written, data are reported as mean with SEM. Regarding comparison between two groups, T-test with Bonferroni correction was chosen and multiple group comparisons was analyzed by using Two-way Anova with multiple comparisons test. P values are shown in figures and are linked to the following meaning; NS non-significant and Significance is specified as *p ≤ 0.05; **p ≤ 0.01, ***p ≤ 0.001.

## SUPPLEMENTAL FIGURE LEGENDS

**Figure S1.**
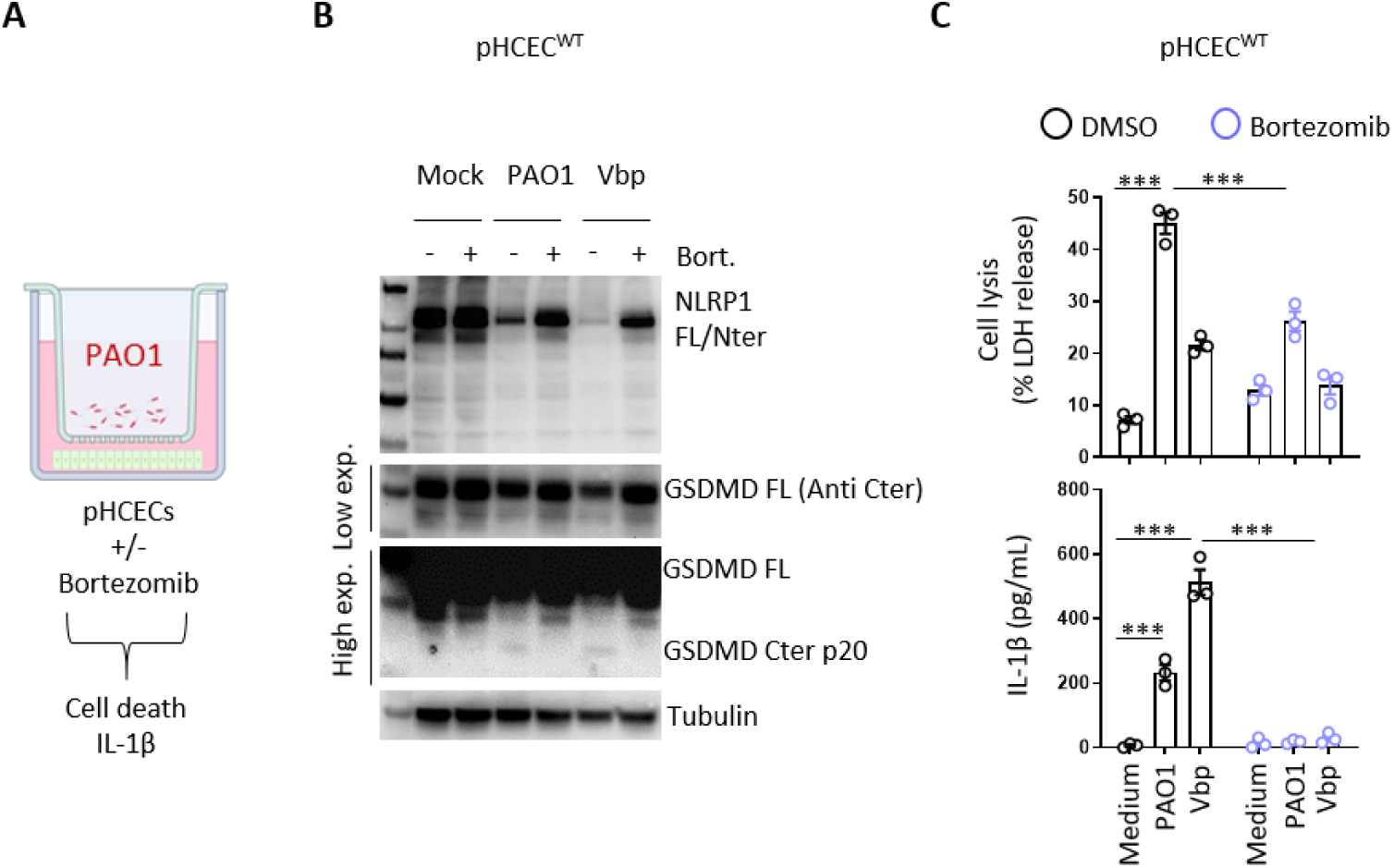
*P. aeruginosa*-activated human NLRP1 inflammasome requires proteasome activity. A. Schematic drawing of *P. aeruginosa* co-culture experiments performed with human corneal epithelial cells. B. Immunoblotting of NLRP1, Gasdermin-D and Tubulin in pHCECs upon Valbororo pro, Vbp 15µM) treatment or *P. aeruginosa* (PAO1, 1.10^5^ bacteria) co-culture for 24 hours in presence/absence of proteasome inhibitor bortezomib. Immunoblots show lysates from one experiment performed at least three times. C. Cell lysis (LDH) and IL-1B release evaluation in pHCECs and pHNECs, upon Valbororo pro, Vbp 15µM) treatment or *P. aeruginosa* (PAO1, 1.10^5^ bacteria) co-culture for 24 hours in presence/absence of proteasome inhibitor bortezomib. ***p ≤ 0.001, Two-Way Anova with multiple comparisons. Values are expressed as mean ± SEM.

**Figure S2.**
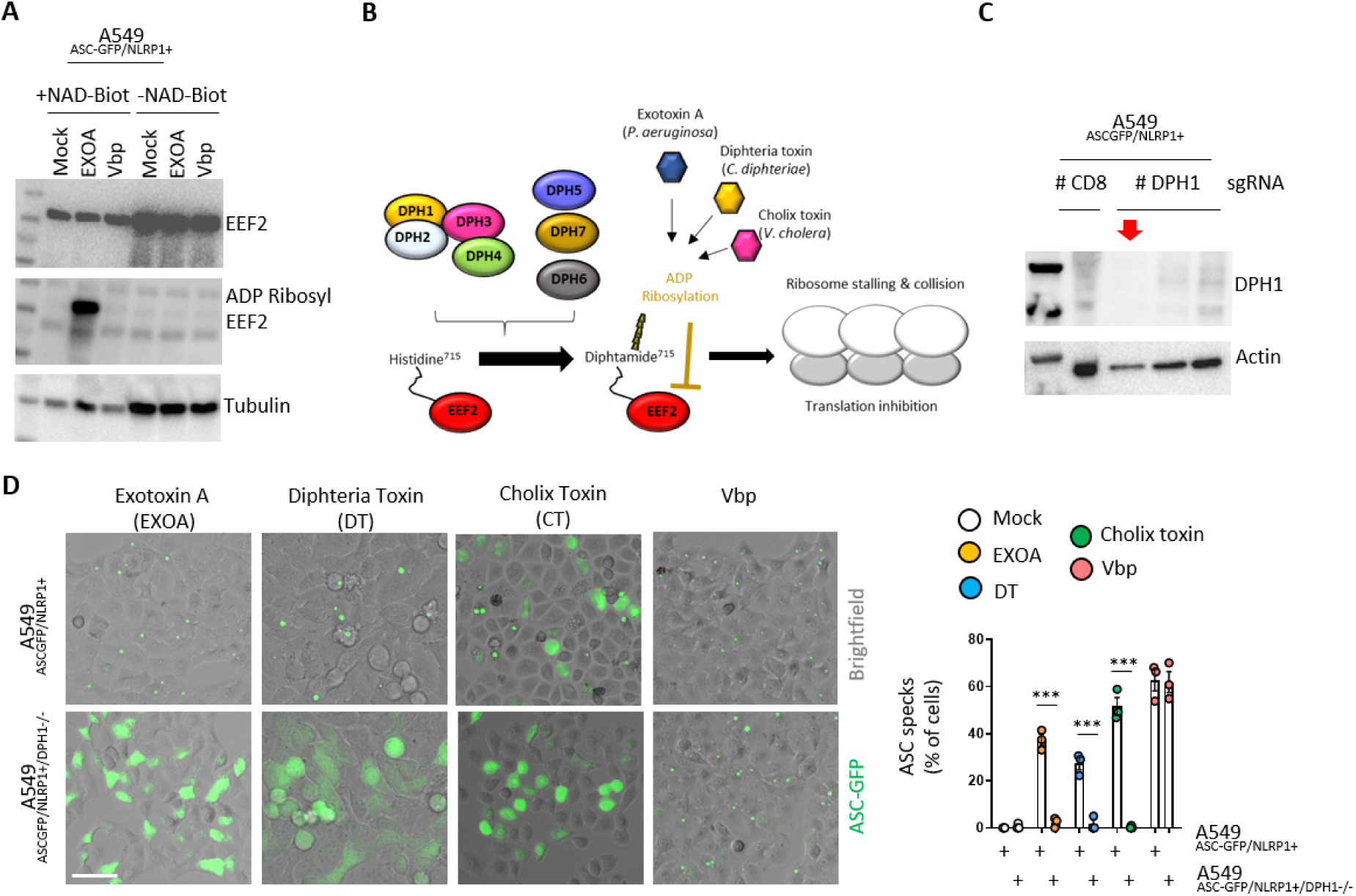
Multiple EEF2-targeting toxins activate the human NLRP1 inflammasome. B. Immunoblotting of ADP-Ribosylated proteins, EEF2 and Tubulin in A549^NLRP1+/ASC-GFP^ cell lysates treated or not with Vbp (15µM) or EXOA (10ng/mL) in presence of NAD-Biot. Immunoblots show lysates from one experiment performed at least three times. B. Schematic mechanism of DPH enzymes at promoting the formation of the Diphtamide aminoacid on EEF2. C. Immunoblotting characterization of genetic invalidation of *DPH1* in A549^NLRP1+/ASC-GFP^ cells using CRISPR-Cas9. D. Fluorescence microscopy and associated quantifications of ASC-GFP specks in A549^NLRP1+/ASC-GFP^ and A549^NLRP1+/ASC-GFP/DPH1-^ reporter cell lines exposed to Vbp (15µM), EXOA (10ng/mL), Cholix Toxin (CT, 10ng/mL) and Diphteria Toxin (DT, 20ng/mL) for 10 hours. ASC-GFP (Green) pictures were taken in dish during after toxin exposure. Images shown are from one experiment and are representative of n = 3 independent experiments; scale bars, 50 µm. ASC complex percentage was performed by determining the ratios of cells positives for ASC speckles (Green, GFP) on the total cells (Birghtfield). At least 10 fields from n = 3 independent experiments were analyzed. Values are expressed as mean ± SEM. ***p ≤ 0.001, One-Way Anova.

**Figure S3.**
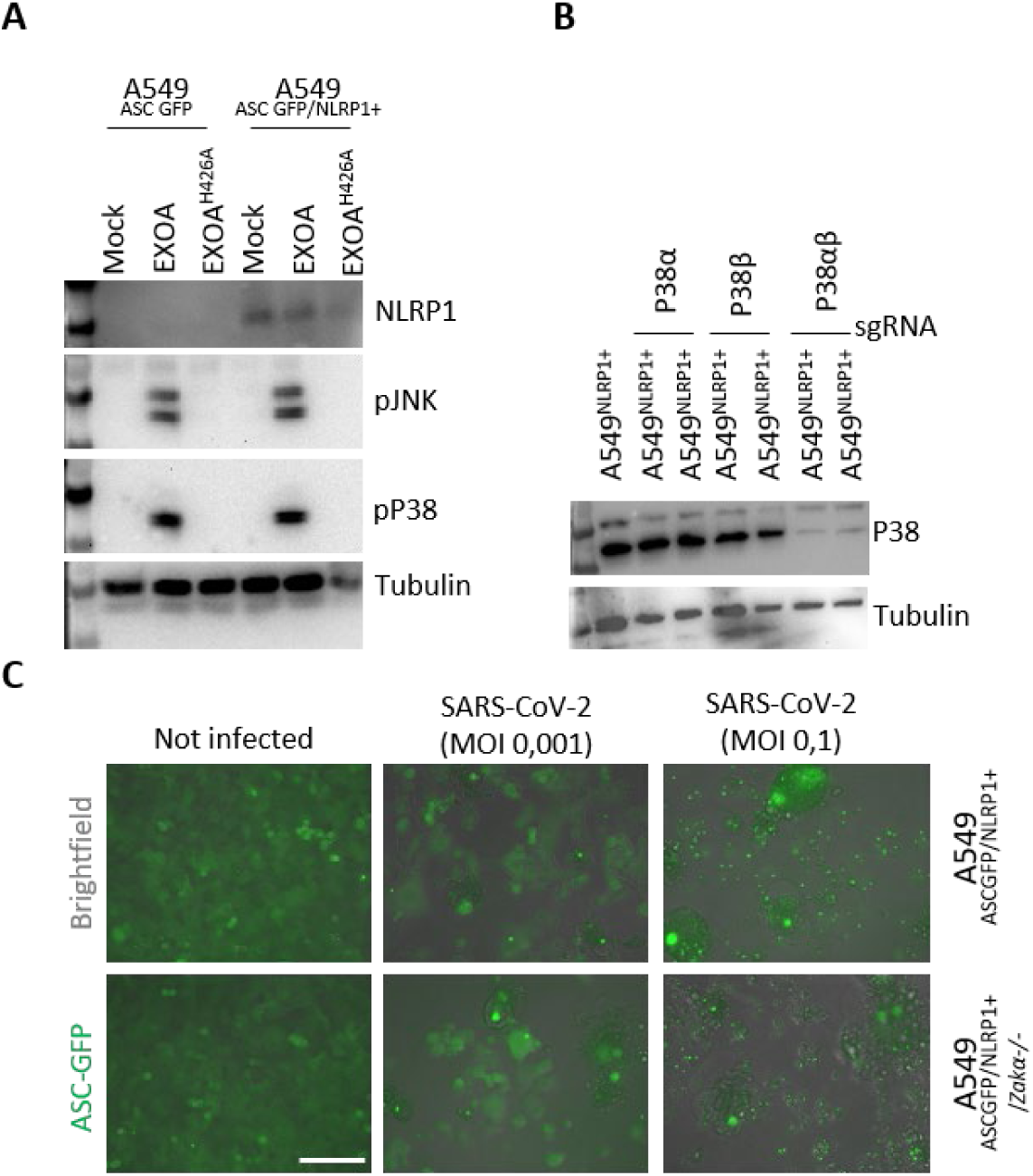
Stress kinase ZAKα regulates EXOA-, but not SARS-CoV-2-driven NLPR1 inflammasome activation. A. Immunoblotting of NLRP1, Tubulin and phophorylated P38 and JNK in A549^NLRP1+^ and A549^NLRP1-^ reporter cell lines exposed or not to EXOA (10ng/mL) or its inactive mutant EXOA^H426A^ for 3 hours. Immunoblots show lysates from one experiment performed at least three times. B. Immunoblotting characterization of genetic invalidation of *P38α and P38β* in A549^NLRP1+/ASC-GFP^ cells using CRISPR-Cas9. C. Fluorescence microscopy and associated quantifications of ASC-GFP specks in A549^NLRP1+/ASC-GFP^ and A549^NLRP1+/ASC-GFP/^*^ZAKα^*^-^ reporter cell lines expressing hACE2 infected for 24 hours with various SARS-CoV-2 MOI (Multiplicity of infection). ASC-GFP (Green) pictures were taken in dish during after toxin exposure. Images shown are from one experiment and are representative of n = 3 independent experiments; scale bars, 50 µm. ASC complex percentage was performed by determining the ratios of cells positives for ASC speckles (Green, GFP) on the total cells (Birghtfield).

**Figure S4.**
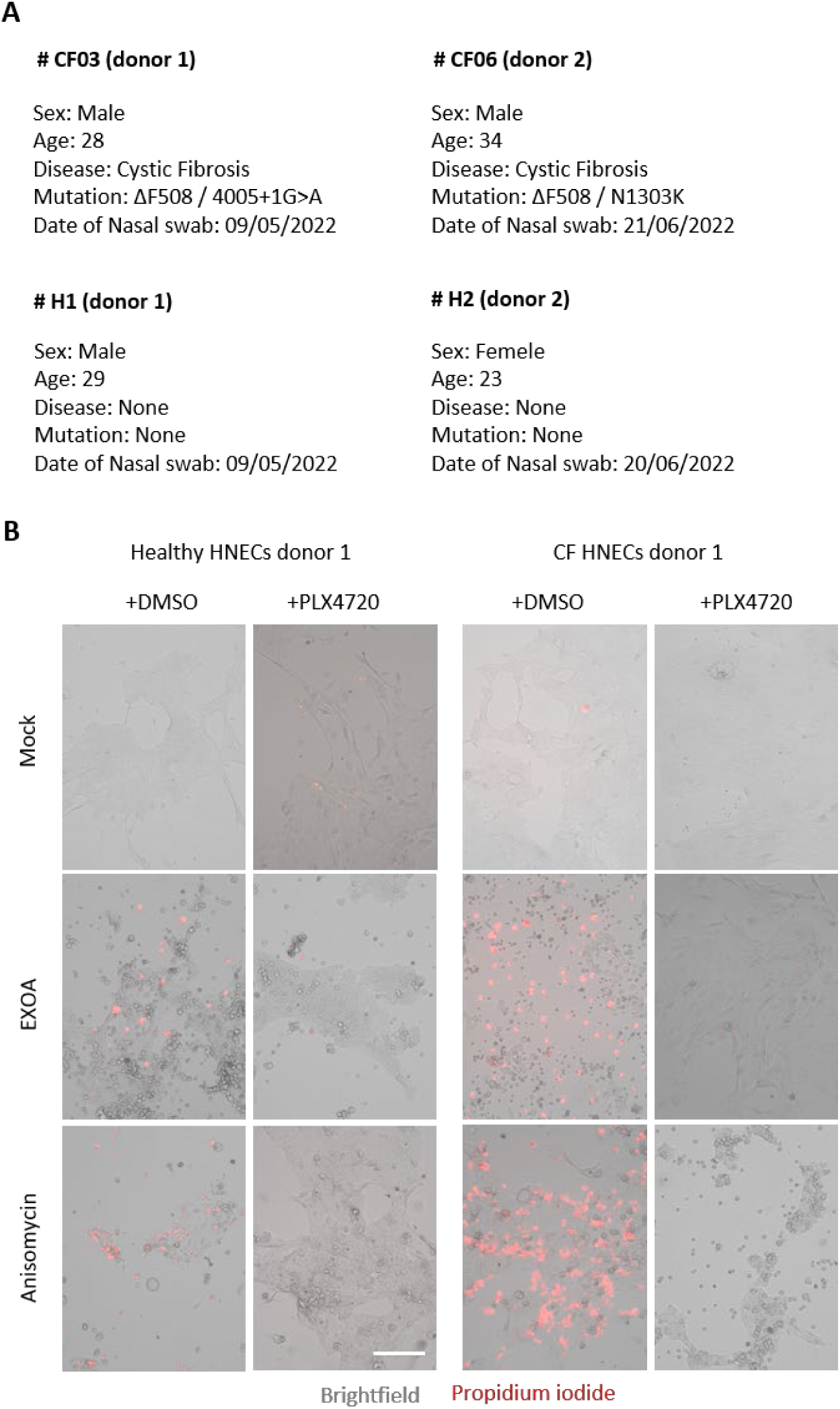
Cystic fibrosis (CF) cells show exacerbated sensitivity to undergo cell death upon ribotoxic stress. A. Information regarding Healthy and CF patient samples used in this study. B. Fluorescence microscopy of Propidium Iodide (PI, red) incorporation into pHNEC^WT^ (donor 2, d2) and pHNEC^CF^ (donor 1, d1) after exposure to Anisomycin or EXOA for 16 hours in presence or not of the ZAK alpha inhibitor PLX4720 (10µM). Images shown are from one experiment and are representative of n = 3 independent experiments; scale bars, 50 µm.

